# Opposing functions of glutamatergic inputs between the globus pallidus external segment and substantia nigra pars reticulata

**DOI:** 10.1101/2023.07.25.550377

**Authors:** Atsushi Yoshida, Okihide Hikosaka

## Abstract

The indirect pathway of the basal ganglia, including the subthalamic nucleus (STN) and globus pallidus external segment (GPe), is believed to play a crucial role in suppressing involuntary movements. However, recent evidence suggests the STN and GPe also facilitate voluntary movements. This study hypothesized that excitatory inputs from the STN to the GPe contribute to this facilitation, and that excitatory projections to the substantia nigra pars reticulata (SNr) are involved in the inhibition. To disrupt the STN-GPe or STN-SNr projections in monkeys during choice and fixation tasks, glutamate receptor inhibitors were injected into the GPe or SNr, which induced delayed saccade latencies toward good choices in the choice task (GPe) and caused frequent reflexive saccades to objects in the fixation task (SNr). Our findings suggest excitatory inputs to the GPe and SNr work in opposing manners, providing new insights that redefine our understanding of the functions of basal ganglia pathways.

**Highlights:**

- STN and GPe neuronal activity increased when good objects were chosen
- SNr activity increased when rejecting bad objects and decreased when accepting good objects
- Excitatory inputs inactivation in the GPe caused delayed saccade to good objects
- Excitatory projection inhibition to the SNr suppressed involuntary saccade control

## Introduction

In the indirect pathway of the basal ganglia circuit, the subthalamic nucleus (STN) and globus pallidus external segment (GPe) are considered to play a vital role in suppressing motor activity,^1–5^ especially involuntary movements. The STN is also implicated in the hyperdirect pathway and is believed to mediate rapid movement inhibition.^6^ However, recent findings have challenged these long-standing beliefs. Studies in both animal models and patients with Parkinson’s disease undergoing deep brain stimulation (DBS) have shown that while a subset of STN neurons display inhibitory activity, other STN neurons modulate excitatory activity that facilitates movement.^7, 8^ Moreover, another study demonstrated that human STN neural code contributes to movement execution.^9^ On the other hand, decreased activity of GPe neurons, which receive direct inhibitory input from the striatum, have been associated with inhibition of inappropriate movement.^10^ However, more than half of all GPe neurons exhibit increased activity, indicating action facilitation.^11–16^ Traditionally, the classic indirect pathway theory was proposed based on anatomical connections from the GPe to the STN.^17–19^ However, as there are many excitatory projections from the STN to the GPe, increased activity in GPe neurons may be caused by these excitatory projections from the STN.^19^ Indeed, electrophysiological experiments have demonstrated that cortical stimulation produces an excitatory response in GPe neurons via the STN.^20, 21^

Therefore, we hypothesized that the STN and GPe play a role in not only suppressing involuntary behavior but also facilitating motor control, potentially through glutamatergic excitatory projections from the STN to the GPe. Experiments in animals with local injections of muscimol, an γ-aminobutyric acid-A (GABA_A_) agonist, are commonly used to investigate the causal relationship between neuronal activity in a certain brain region and its behavior.^22^ Inactivation by muscimol allows us to examine whether the region itself is involved in a certain function, but not which pathways are involved. Previous studies have used glutamate receptor antagonists to examine how excitatory projections affect the neural activity of the GPe,^21, 23^ but their concentrations were too low to induce behavioral changes Thus, we assumed that high concentrations of glutamate receptor antagonists would produce behavioral changes and allow us to examine the function of excitatory projection from the STN.

To test our hypothesis, we first identified the task-related regions in the STN and GPe by recording the neural activity while two monkeys performed a motivational choice task in which they either accepted a good object that yielded a liquid reward or refused a bad object that yielded no reward. Thereafter, we evaluated changes in their behavior during the task after injecting a glutamate receptor antagonist into the GPe to perturb STN-GPe projections. For comparison, we also injected glutamate receptor antagonists in the substantia nigra pars reticulata (SNr) to inhibit the STN-SNr projection, which has been implicated in both indirect and hyperdirect pathways. This study reveals that local injection of glutamatergic antagonists into the GPe delayed saccade reaction times toward contralateral good objects, whereas injection into the SNr accelerated saccade reaction times to contralateral bad objects. Our results suggest the opposite functions of excitatory projections to GPe or to SNr.

## Results

### Behavior results in the choice task

Two monkeys performed the choice task, as illustrated in Figure 1A. In this task, one of the four *scenes* from a chosen set was randomly displayed (Figure 1B). Each *scene* suggested that either a good object, leading to reward delivery, or a bad object, yielding no reward, would be presented sequentially and randomly. In cases where the monkeys saccaded toward the object and maintained fixation, it was interpreted as object acceptance. If the monkeys initiated a saccade but did not fixate on it and returned to the center point (*return*) or persistently gazed around the center point (*stay*), these actions were categorized as object rejection (Figure 1C); after which, the object disappeared and another object was presented. This sequence was repeated until the monkey accepted an object. In *scenes* 3 and 4, the values of the paired objects alternated (Obj E was good and Obj F was bad in *scene* 3, while Obj E was bad and Obj F was good in *scene* 4). *Scenes* 3 and 4 were used to examine whether the neuronal responses were associated with object values or visual characteristics. For individual neuronal recordings, one of the six available sets was randomly chosen (Figure S1).

**Figure 1.**
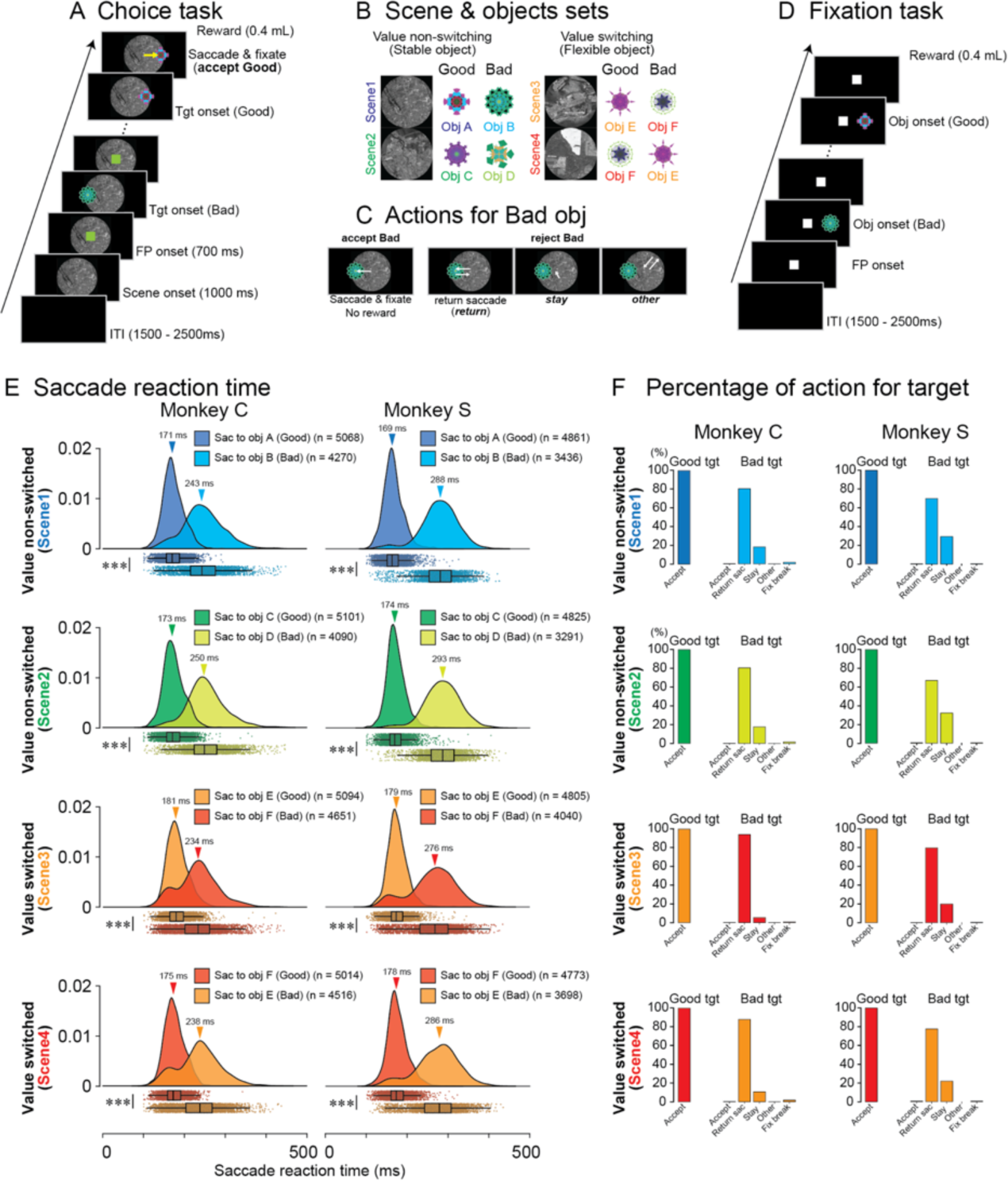
Task paradigms and Behavioral results. (A, B) Task procedure of the choice task (A) and one example set for recording from a single neuron (B). One set included four *scenes*, and each *scene* was combined with one good (rewarded) and one bad object (non-rewarded) in the choice task. In *scenes* 1 and 2, object values are stable, while in *scenes* 3 and 4, the same two objects were used, but their values were switched (flexible) in each *scene*. (C) Chosen actions toward bad objects in the choice task. Monkeys rejected bad objects by making a return saccade (*return*), looking around the center point (*stay*), or making a saccade away from bad objects (*other*). (D) Task procedure of the fixation task. Monkeys were required to suppress a saccade toward presented objects. Objects used in the fixation task were the same as objects in *scene* 1 in the choice task. (E) Raincloud plots of saccade reaction times toward good or bad objects in *scenes* 1-4 during the choice task of two monkeys, C and S. In each panel, the upper “cloud” shows the probability distribution of reaction times, and the lower “rain” illustrates the raw plots of individual reaction times. In all *Scenes* of the two monkeys, the reaction times toward good objects were significantly shorter than those toward bad objects (*p* = 0 for all *scenes* in both monkeys, Welch’s t-tests). (F) Percentages of selected actions for good and bad objects in *scenes* 1-4 during the choice task.

Figure 1E shows the raincloud plots of the saccade reaction times for good (saccade for accept) or bad objects (*return* saccade for reject) in each scene for both monkeys. The “cloud” represents the data distributions, while the “rain” shows jittered raw data. Boxplots, including median values and confidence intervals, were overlaid on the rain data plots.^24^ Table 1 summarizes the number of samples, mean, standard deviation (SD), and 95% confidence intervals (CI) of the reaction times toward good or bad objects in each *scene*, collected during neuronal recording sessions. Welch’s t-test was employed to compare the reaction times toward good and bad objects in *scenes* 1–4 and revealed significant differences in each *scene*. These findings suggest that monkeys recognize objects as good or bad when they make a saccade toward them.

**Table 1.** Saccade reaction times in each condition of each monkey.

| **Monkey C** | n | mean RT (ms) | SD | 95% CI |
| --- | --- | --- | --- | --- |
| **good obj** |  |  |  |  |
| ObjA in scene1 | 5068 | 174.6 | 26.3 | [173.9 175.3] |
| ObjC in scene2 | 5101 | 175.9 | 26.7 | [175.2 176.7] |
| ObjE in scene3 | 5094 | 184.2 | 27.1 | [183.5 185.0] |
| ObjF in scene4 | 5014 | 177.6 | 25.1 | [176.9 178.3] |
| **bad obj** |  |  |  |  |
| ObjB in scene1 | 4270 | 245.9 | 49.6 | [244.5 247.4] |
| ObjD in scene2 | 4090 | 254.3 | 47.4 | [252.9 255.8] |
| ObjF in scene3 | 4651 | 232.7 | 50.1 | [231.3 234.2] |
| ObjE in scene4 | 4516 | 236.8 | 51.8 | [235.3 238.4] |
| **Monkey S** | n | mean RT (ms) | SD | 95% CI |
| **good obj** |  |  |  |  |
| ObjA in scene1 | 4861 | 171.4 | 26.2 | [170.7 172.2] |
| ObjC in scene2 | 4825 | 176.9 | 25.8 | [176.2 177.6] |
| ObjE in scene3 | 4805 | 177.1 | 26.2 | [180.3 181.8] |
| ObjF in scene4 | 4773 | 177.0 | 27.2 | [180.1 181.7] |
| **bad obj** |  |  |  |  |
| ObjB in scene1 | 3436 | 286.3 | 47.3 | [284.8 288.0] |
| ObjD in scene2 | 3291 | 292.1 | 46.1 | [290.6 293.8] |
| ObjF in scene3 | 4040 | 269.7 | 57.8 | [268.0 271.5] |
| ObjE in scene4 | 3698 | 280.7 | 55.0 | [279.0 282.5] |

Figure 1F shows the proportion of actions chosen by the monkeys for the good and bad objects. Table 2 shows the number of actions directed toward bad objects in each *scene*. We found that the monkeys consistently accepted good objects, whereas for bad objects, monkeys frequently made a *return* (monkey C: 80.6%, 80.5%, 93.5%, and 87.3% in *scenes* 1–4, respectively; monkey S: 69.4%, 66.7%, 78.7%, and 77.2% in *scenes* 1–4, respectively) and *stay* (monkey C: 17.3%, 17.3%, 5.3%, and 10.4% in *scenes* 1–4, respectively; monkey S: 29.8%, 32.4%, 20.7%, and 21.9% in *scenes* 1–4, respectively). During the task, the target range was defined as an 8°-per-side square; when monkeys executed a saccade to the target but their eye position exited the range for <400 ms, the target vanished, and the fixation point reappeared. In other words, the monkeys had to wait 400 ms for the next target to be presented if they rejected bad objects by *stay*, while the waiting time for the next target was shorter than 400 ms if they rejected bad objects by *return*, which may explain the frequent rejection of bad objects by *return*. Comparing the percentages of *stay* among the four scenes, the percentages of *stay* rejections in *scenes* 1 and 2 for both monkeys were significantly higher than those in *scenes* 3 and 4 (Fisher’s test, monkey C; *p* < 0.0001, φ = 0.15, 95% CI: 0.37–0.44; monkey S: *p* < 0.0001, φ = 0.11, 95% CI: 0.56–0.64). This result may be attributed to the consistent object values in *scenes* 1 and 2, and the alternating values of the objects in *scenes* 3 and 4.

**Table 2.** Counts of chosen actions for bad objects.

| **Monkey C** | total | accept | return | stay | other | fxbreak |
| --- | --- | --- | --- | --- | --- | --- |
| scene1 | 5367 | 12 | 4326 | 932 | 6 | 91 |
| scene2 | 5166 | 9 | 4157 | 898 | 2 | 100 |
| scene3 | 5054 | 22 | 4724 | 267 | 3 | 38 |
| scene4 | 5239 | 25 | 4574 | 545 | 2 | 93 |
| non-switch(scene1,2) | 10533 | 21 | 8483 | 1830 | 8 | 191 |
| switch(scene3,4) | 10293 | 47 | 9298 | 812 | 5 | 131 |
| **Monkey S** | total | accept | return | stay | other | fxbreak |
| scene1 | 4877 | 17 | 3383 | 1455 | 0 | 22 |
| scene2 | 4862 | 25 | 3242 | 1575 | 0 | 20 |
| scene3 | 5081 | 12 | 4001 | 1051 | 0 | 17 |
| scene4 | 4751 | 19 | 3669 | 1042 | 0 | 21 |
| non-switch(scene1,2) | 9739 | 42 | 6625 | 3030 | 0 | 42 |
| switch(scene3,4) | 9832 | 31 | 7670 | 2093 | 0 | 38 |

### STN, GPe, and SNr neuronal activity during the choice and fixation tasks

To explore the involvement of the STN, GPe, and SNr neurons in the acceptance or rejection of objects, we recorded neuronal activity in these three areas while the monkeys performed the choice task. We identified two types of task-related neurons in the STN and the GPe (Figures 2A, D, G, and J). One group of neurons demonstrated increased activity when choosing good objects compared to the activity seen when rejecting bad objects and were defined as good-preferring (Good-pref) STN (n = 151; 81 from monkey C and 70 from monkey S) and GPe (n = 124; 65 from monkey C and 59 from monkey S) neurons (Figures 2A and G). The other group of STN neurons exhibited increased activity during the rejection of bad objects, whereas the other type of GPe neurons showed a greater decrease in activity, and were defined as bad-preferring (Bad-pref) STN (n = 32; 10 from monkey C, 22 from monkey S) and GPe (n = 81; 46 from monkey C, 35 from monkey S) neurons (Figures 2D and J). Conversely, task-related SNr neurons (n = 100; 51 from monkey C, 49 from monkey S) exhibited strong activity modulation, whereby both the acceptance of good objects decreased activity and rejection of bad objects increased activity (Figure 2M). The number of recorded neurons and specific recording sites in the STN, GPe, and SNr are summarized in Table 3 and Figure S2, respectively.

**Figure 2.**
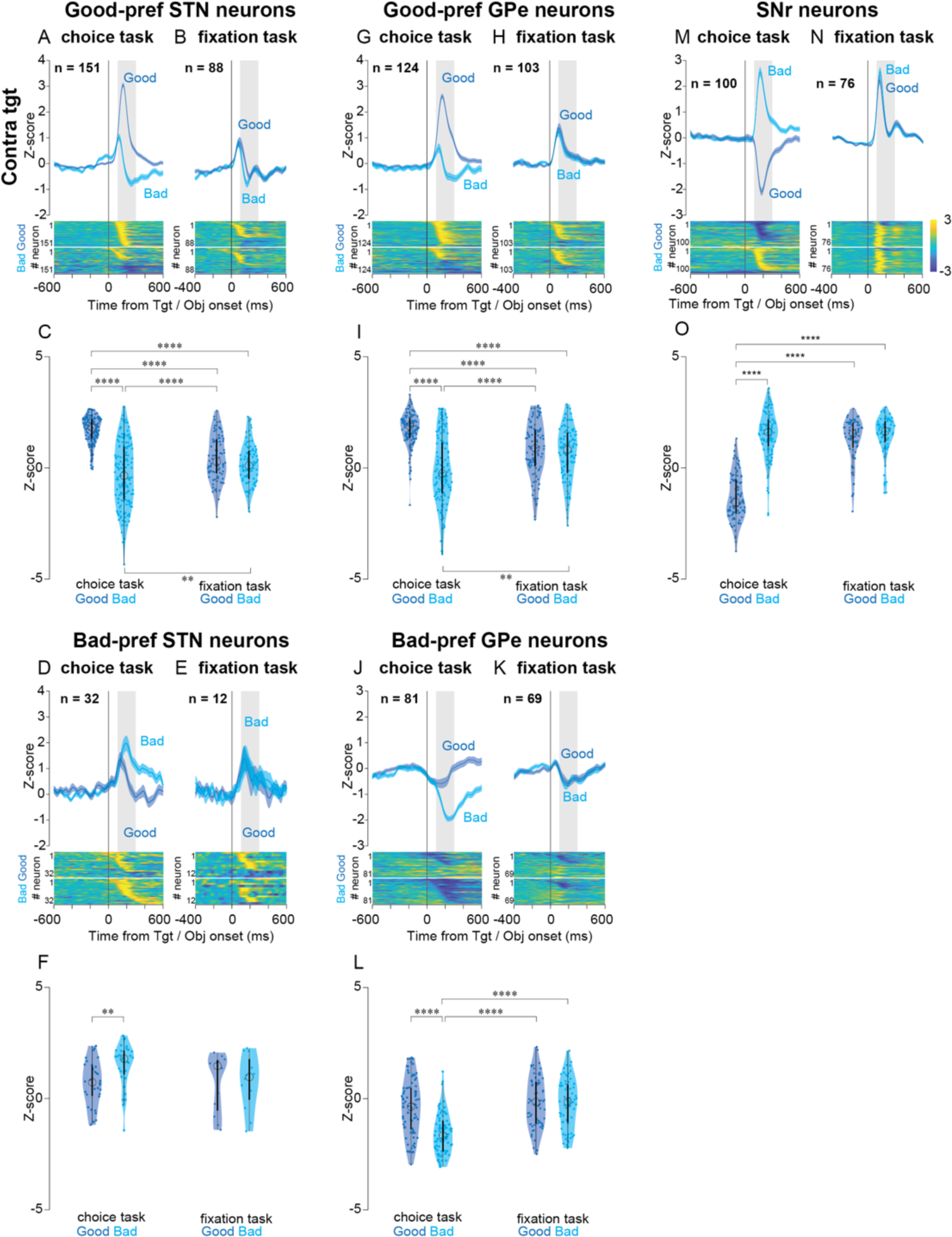
Population activity of Good-& Bad-pref STN, Good-& Bad-pref GPe, and SNr neurons at contralateral target onset in the choice and fixation task. (A, D, G, J, and M) Normalized mean population activity aligned at contralateral good and bad target onset and the color maps of the normalized firing rate of individual neurons during the Choice task of Good-pref STN neurons (A), Bad-pref STN neurons (D), Good-pref GPe neurons (G), Bad-pref GPe neurons (J), and SNr neurons (M). Black-colored lines demonstrate the population activity when monkeys rejected bad objects by *stay* (looking around the center point). (B, E, H, K, and N) Normalized mean population activity aligned at contralateral good and bad object onset and the color maps of the normalized firing rate of individual neurons during the fixation task of Good-pref STN neurons (B), Bad-pref STN neurons (E), Good-pref GPe neurons (H), Bad-pref GPe neurons (K), and SNr neurons (N). (C, F, I, L, and O) The violone plots of the mean normalized firing rate of individual neurons during target onset in the choice task and object onset in the fixation task of Good-pref STN neurons (C), Bad-pref STN neurons (F), Good-pref GPe neurons (I), Bad-pref GPe neurons (L), and SNr neurons (O). Neuronal activity was measured for a 200-ms interval 100 ms after target onset (gray rectangle in (A, D, G, J, and M)) and object onset (gray rectangle in (B, E, H, K, and N)). The asterisk indicates a significant difference in normalized neuronal activity in comparison among the conditions in the choice and fixation tasks (post-hoc pairwise t-tests with Bonferroni correction, \**p <* 0.05, \*\**p* < 0.01, \*\*\**p* < 0.001, \*\*\*\**p* < 0.0001). Asterisks are attached only to combinations with significant differences.

**Table 3.** Numbers of recorded task-related neurons in the STN, GPe, and SNr of each monkey.

|  | STN |  | GPe |  | SNr |
| --- | --- | --- | --- | --- | --- |
|  | Good-pref | Bad-pref | Good-pref | Bad-pref |  |
| **choice task** |  |  |  |  |  |
| Monkey C | 81 | 10 | 65 | 46 | 51 |
| Monkey S | 70 | 22 | 59 | 35 | 49 |
| total | 151 | 32 | 124 | 81 | 100 |
| **fixation task** |  |  |  |  |  |
| Monkey C | 65 | 9 | 49 | 37 | 41 |
| Monkey S | 21 | 3 | 53 | 35 | 32 |
| total | 88 | 12 | 102 | 69 | 73 |

To determine whether the neuronal activities of the STN, GPe, and SNr neurons observed in the choice task were associated with the suppression of involuntary saccades to presented objects, we used a fixation task for a subset of neurons that modulated activity in the choice task (Figure 1D). No distinct differences in STN, GPe, or SNr neuronal activity between good and bad objects were observed in the fixation task (Figures 2B, E, H, K, and N). Although the degree of neuronal modulation of Good-pref STN neurons and Good- and Bad-pref GPe neurons observed in the choice task was reduced in the fixation task (Figures 2B, H, and K), Bad-pref STN and SNr neurons exhibited comparable activity modulation for bad objects in both the choice and fixation tasks (Figures 2E, and N). Notably, the neuronal activity of SNr neurons for good objects in the fixation task increased, showing a similar response to that of bad objects in the choice and fixation tasks (Figure 2N). These findings suggest that the increased activity for good objects observed in Good-pref STN and GPe neurons in the choice task may facilitate saccades for selecting good objects, and that the increased activity for bad objects in SNr neurons during the choice task may be related to the suppression of involuntary saccades.

To analyze the data quantitatively, we employed a parametric bootstrap test for linear mixed-effect models (LMMs) and subsequent post-hoc pairwise t-tests with Bonferroni correction (see Methods for details). The violin plots in Figures 2C, F, I, L, and O depict the mean normalized firing rate of individual STN, GPe, and SNr neurons during the 200-ms period from 100–300 ms following target onset in both the choice and fixation tasks. Statistical analysis revealed a significant difference in neuronal activity between good and bad objects in the choice task (post-hoc pairwise t-tests, *p* < 0.0001 for Good-pref STN, Good-pref GPe, Bad-pref GPe, and SNr neurons, and *p* < 0.01 for Bad-pref STN neurons), whereas no significant difference in activity was found between good and bad objects in the fixation task. The detailed results of the other post-hoc pairwise test comparisons are summarized in Table S1.

To assess whether the neuronal activity observed in the choice task reflected the visual characteristics of the presented objects or their values, we utilized a set of four conditions (*scenes* 1–4). Neurons in the STN, GPe, and SNr modulated their activity upon *scene* presentation (Figure S3), with no significant differences among *scenes* 1–4 (parametric bootstrap tests, full model vs. null models; *p* = 0.06, Good-pref STN; *p* = 0.68, Bad-pref STN; *p* = 0.54, Good-pref GPe; *p* = 0.13, Bad-pref GPe; *p* = 0.66, SNr). Significant differences were observed within *scenes* 1–4 for the normalized neuronal activity in the STN, GPe, and SNr during the 200-ms period from 100 ms after the target onset in the choice task (Figures S4–S6). The detailed results of the post-hoc pairwise test comparisons are summarized in Tables S2–S6. We also conducted statistical tests using the data aligned with saccade initiation (Figures S7–S9). Although there were significant differences in some conditions (Good-pref STN, contralateral Bad between *scenes* 1 and 3 (*p* < 0.01), contralateral Bad between *scenes* 2 and 3 (*p* < 0.0001), contralateral Bad between *scenes* 2 and 4 (*p* < 0.001); Good-pref GPe, contralateral Bad between *scenes* 2 and 3 (*p* < 0.01)), most conditions across *scenes* to 1–4 were not significantly different. The detailed results of the other post-hoc pairwise test comparisons are summarized in Tables S7–S11.

### Opposite effects of glutamatergic antagonist on the GPe and SNr

Based on the results of the activity of the STN, GPe, and SNr neurons during the choice and fixation tasks, as well as previous anatomical and electrophysiological studies,^19–21^ we hypothesized that Good-pref STN neurons send excitatory projections to Good-pref GPe neurons, which, in turn, cause a decrease in the firing rates in SNr neurons, thereby facilitating a saccade toward good objects in the choice task; conversely, Bad-pref STN neurons may have excitatory projections to SNr neurons to suppress saccades during the choice and fixation tasks. In other words, glutamatergic excitatory projections from the STN to the GPe may facilitate voluntary saccades, while the excitatory projections from the STN to the SNr may be related to the inhibition of involuntary saccades.

To test this hypothesis, we locally injected a mixture of N-methyl-d-aspartate (NMDA) receptor antagonist (carboxypiperazin-4-propyl-1-phosphonic acid, CPP) and aminomethylphosphonic acid (AMPA) receptor antagonist (2,3-dihydroxy-6-nitro-7-sulfamoyl-benzo (F)quinoxaline, NBQX) into the GPe or SNr, where many task-related neurons were found when the monkeys performed the choice and fixation tasks. Saccade reaction times toward contralateral good objects in the choice task were significantly delayed following glutamatergic antagonist injection into the caudal-dorsal GPe (post-hoc pairwise t-tests, *p* < 0.0001, Figure 3A). Moreover, the proportion of *return* for bad objects in both directions decreased significantly (*p* < 0.0001, for both directions, Figure S10A), while the proportion of *stay* increased in both directions (*p* < 0.001, for contralateral, and *p* < 0.0001, for ipsilateral, Figure S10A). By contrast, glutamatergic antagonist injection into the lateral SNr led to faster saccade reaction times for good and bad contralateral objects in the choice task (*p* < 0.001, for good objects, and *p* < 0.0001, for bad objects, Figure 3B). Additionally, the proportion of *return* toward contralateral bad objects significantly increased (*p* < 0.0001, for both directions, Figure S10B), whereas the proportion of *return* toward ipsilateral bad objects significantly decreased (*p* < 0.0001, for both directions, Figure S10B). In the fixation task, the proportion of fixation break errors significantly increased when objects were presented on the contralateral side after glutamatergic antagonist injection into the SNr (*p* < 0.0001, Figure 3D), but no significant changes were observed in the GPe. Control experiments using saline showed no significant changes in behavior for either task. These findings suggest that excitatory input, which may be mainly from the STN, to the GPe facilitates voluntary saccades, whereas inputs from the STN to the SNr suppress involuntary saccades. The detailed results of the statistical test comparisons are summarized in Tables S12– S15.

**Figure 3.**
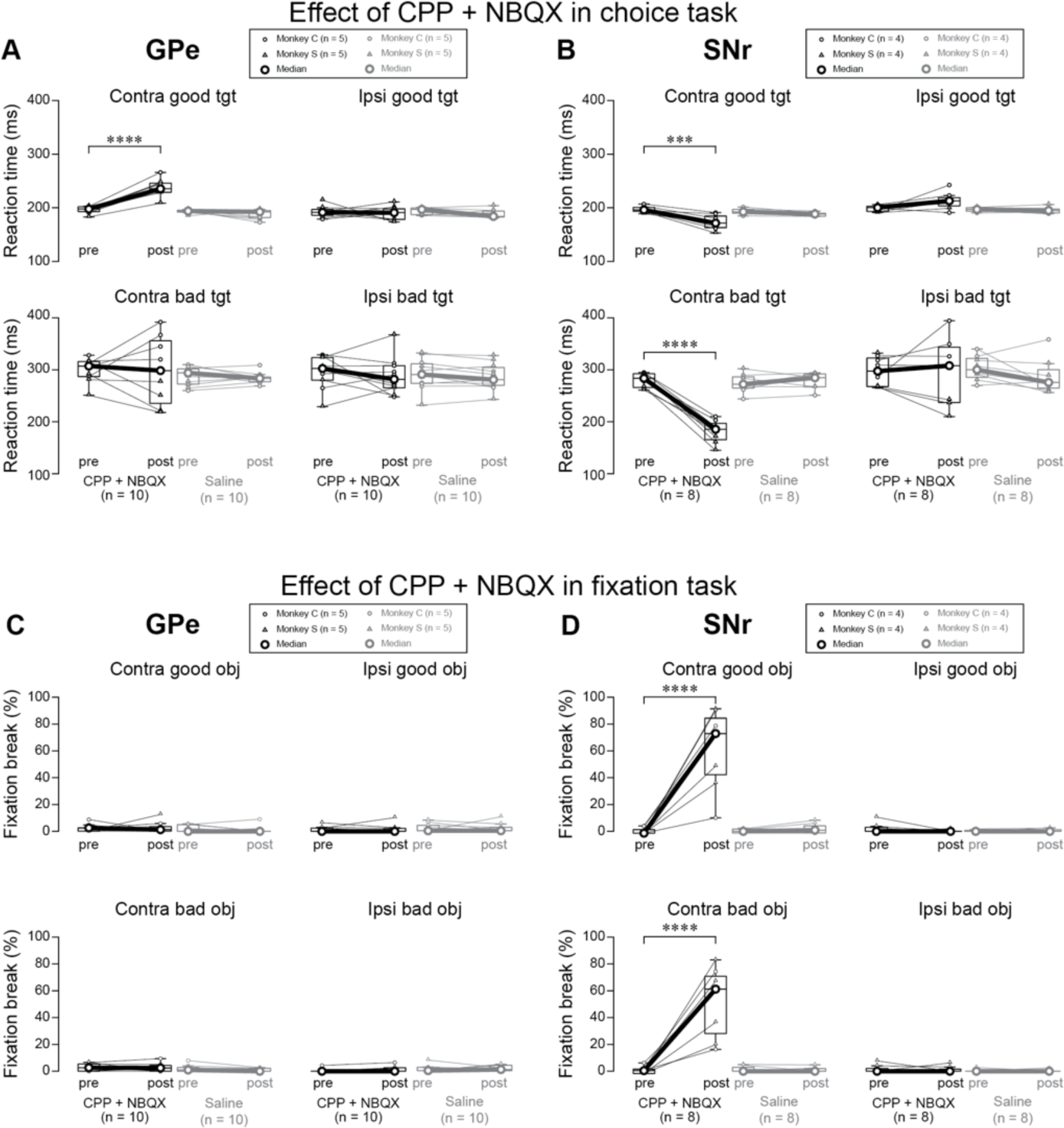
Effects of glutamatergic antagonist injection into GPe and SNr during the choice task and fixation task. (A, B) Changes of saccade reaction time toward good and bad targets in the choice task before and after injection of CPP (NMDA-type receptor antagonist) and NBQX (AMPA-type receptor antagonist) mixture or saline into the GPe (A) and SNr (B). The asterisk indicates a significant difference in median saccade reaction times in comparison among the conditions in the choice tasks (post-hoc pairwise t-tests with Bonferroni correction, \**p <* 0.05, \*\**p* < 0.01, \*\*\**p* < 0.001, \*\*\*\**p* < 0.0001). Asterisks are attached only to combinations with significant differences. (C, D) Comparison of fixation break rates in the fixation task when CPP + NBQX or saline were injected into the GPe (C) and SNr (D). The asterisk indicates a significant difference in the median proportion of fixation break error in comparison among the conditions in the fixation tasks (post-hoc pairwise t-tests with Bonferroni correction, \**p <* 0.05, \*\**p* < 0.01, \*\*\**p* < 0.001, \*\*\*\**p* < 0.0001). Asterisks are attached only to combinations with significant differences.

### Injection sites in the GPe and SNr visualized using quantitative susceptibility mapping (QSM)

We next applied QSM during magnetic resonance imaging (MRI) to visualize the structures of the basal ganglia and detect altered regions of magnetic susceptibility by quantitatively analyzing iron deposition (Figure 4A).^25^ Compared to conventional T1- and T2-weighted images, QSM could successfully visualize basal ganglia structures. Notably, the lateral side of the SNr, where many task-related neurons were found, exhibited a higher QSM signal than that observed in other SNr regions (Figure 4B) and was thought to correspond with the substantia nigra pars lateralis (SNpl).^26–29^

**Figure 4.**
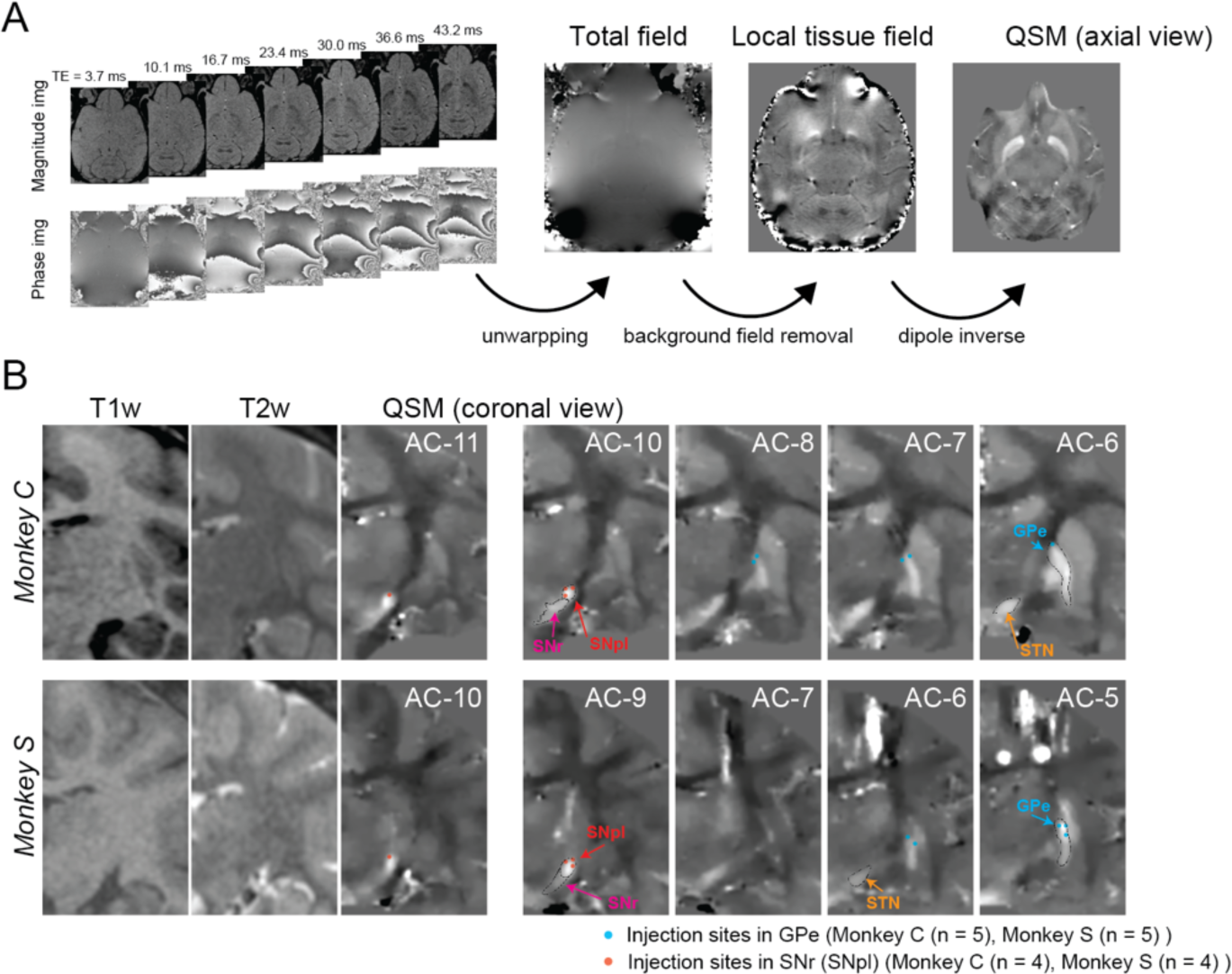
Injection sites on images of quantitative susceptibility mapping (QSM) and the new concept of the basal ganglia circuit. (A) Overview of QSM processing. QSM image is reconstructed from phase images with different TE of 3D gradient echo sequence. In the processing from the raw phase image to QSM, there are three steps, unwarping, background phase removal by high-pass filtering, and inverse problem solutions by dipole deconvolution. (B) Plots of injection sites into the GPe and SNr on QSM images. Abbreviations: AC, anterior commissure. GPe, globus pallidus external segment. SNpl, substantia nigra pars lateralis. SNr, substantia nigra pars reticulata. STN, subthalamic nucleus.

## Discussion

For decades, it has been assumed that the STN and GPe are involved in the suppression of involuntary movement because of their role in the indirect pathway of the basal ganglia. However, several studies have shown that STN and GPe neuronal activity changes when subjects intentionally choose to perform an action. In the present study, many STN and GPe neurons strongly increased their activity when the monkeys made saccades toward good objects (Good-pref STN and GPe neurons) during the choice task, while the degree of neuronal modulation during the choice task in the STN and GPe decreased during the fixation task. Moreover, saccade reaction times were delayed for good contralateral objects in the choice task when a glutamatergic antagonist was injected into the GPe. Additionally, saccade latencies became shorter for contralateral objects in the choice task when the glutamatergic antagonist was injected into the SNr, which also induced frequent fixation break errors to the contralateral side in the fixation task.

### STN and GPe neurons are also involved in action facilitation

Contrary to the conventional belief that STN and GPe neurons are involved in the suppression of inappropriate motor activity, we found many Good-pref neurons in the STN and GPe that significantly altered neuronal activity when choosing a good object. In particular, Good-pref neurons comprised the majority of neurons in the STN, consistent with the findings of previous studies.^7, 8^ These results suggest that the STN and GPe are involved in the inhibition and facilitation of involuntary and appropriate motor activity.

In our previous studies, we found saccade-related neurons in the ventral part of the STN^30, 31^ and projections from the frontal and supplementary eye fields to the ventral STN,^32, 33^ consistent with the presence of many task-related neurons in the ventral STN (Figure S2A) as identified in the current study. We also noted many task-related neurons in the caudal-dorsal part of the GPe (cdGPe) (Figure S2B). Although we did not investigate evidence of direct connections between the task-related STN and GPe neurons in this study, a previous anatomical study in macaque monkeys illustrated abundant connections between the ventral STN and dorsal GPe.^19^ Therefore, the results of finding task-related neurons in the ventral STN and dorsal GPe are anatomically reasonable.

Another previous study of ours indicated that caudal-ventral GPe (cvGPe) neurons receive numerous projections from the tail of the caudate nucleus,^34^ while neuronal activity in the cvGPe showed decreased activity when bad values of objects were presented in the passive-viewing task, which is comparable to the fixation task in the present study; the neuronal modulation in the cvGPe was considered to be related to visual response. In the current study, we showed that the degree of neuronal modulation in the cdGPe was greater in the choice task than in the fixation task, suggesting that the function of cdGPe neurons is motor-related, while that of cvGPe neurons is visual-related.

### Intensive activation of SNr neurons in the fixation task

The intensive neuronal modulation of Good-pref STN and GPe neurons during the acceptance of good objects observed in the choice task was diminished in the fixation task, indicating that the activity of these neurons in the choice task was reflected in saccade initiation to choose a good object. By contrast, the degree of modulation of SNr neurons in the fixation task was comparable to that of the rejection of bad objects in the choice task. Importantly, SNr neurons decreased their activity when the monkeys selected a good object in the choice task, but increased activity when good objects were presented in the fixation task, indicating that this increased activity in the fixation task may be related to the suppression of saccades toward objects. Thus, this excitatory activity of SNr neurons in the choice and fixation tasks may reinforce inhibitory projections to the superior colliculus and involve the suppression of saccade initiation.

### Opposite effects of glutamatergic input perturbation on the GPe and SNr

Local injection of glutamatergic antagonists, CPP and NBQX, into the GPe induced a significant delay in saccade reaction times to the contralateral good object during the choice task, suggesting that glutamatergic excitatory projections to the GPe are involved in facilitating saccade generation. Previous anatomical studies have shown that excitatory projections to the GPe are mainly from the STN^19^ and that the STN-GPe pathway is related to saccade facilitation.

For comparison, we locally injected the glutamatergic antagonist into the lateral part of the SNr, where we found numerous task-related neurons. After administration, saccade reaction times to the contralateral objects in the choice task became shorter, with frequently occurring fixation break errors in saccades to the contralateral objects. These results suggest that the monkeys were unable to suppress reflexive saccades toward contralateral objects, suggesting that glutamatergic input to the SNr is involved in the suppression of reflexive actions. Therefore, in addition to the excitatory projections to the GPe, it is likely that the STN-SNr pathway mediates the inhibition of reflexive actions since the projections to the SNr are thought to be mainly from the STN.^35, 36^

### STN-GPe pathway explains the action facilitation by DBS

Previous studies have provided explanations based on the conventional concept of basal ganglia pathways, which have been widely accepted to explain pathological conditions such as Parkinson’s disease. However, certain phenomena cannot be explained using this conventional concept. Indeed, DBS of the STN in Parkinson’s disease has been shown to improve not only the inhibition but also the facilitation of motor function.^37–39^ However, if the STN-GPe pathway reported in the present study is improved by DBS, the effect of DBS on STN neuronal activity may be mediated not only via the STN-SNr pathway (behavioral inhibition) but also through the STN-GPe pathway (appropriate action facilitation). Therefore, the inclusion of our proposed STN-GPe pathway to the existing theory may lead to a better understanding of normal basal ganglia function and basal ganglia-related disease pathophysiology.

### Revisit of the SNpl

The SNr has been implicated in the involvement of saccade generation. However, a closer look at the recording sites of saccade-related neurons of the SNr in previous studies revealed many neurons mainly in the lateral part of the SNr, as in the present study.^40–42^ This site is called the SNpl, and previous anatomical studies in monkeys in the 1980s have indicated that it differed from the SNr in terms of connectional, cytoarchitectural, and histochemical aspects.^27, 28, 43, 44^ Anatomical studies have also illustrated that neurons in the SNpl send efferent projections mainly to the superior colliculus, the center of eye movement in the brainstem,^26, 27, 45, 46^ while they receive afferent projections from the dorsal part of the GPe,^47^ where many saccade-related neurons were found in the present and our previous studies.^13, 14^ Although we did not reveal no direct anatomical connections between task-related neurons in the GPe and SNr neurons in this study, our results are consistent with the neurophysiological and anatomical findings of previous studies. Additionally, we used a new MRI method called QSM and found that the QSM signals were higher in the lateral part of the SNr than in the residual SNr parts; this region probably coincides with the SNpl. Thus, QSM may be useful in identifying the SNpl.

### Limitations

The current study has some limitations that warrant discussion. First, the possibility that excitatory projections from regions other than the STN to the GPe cannot be ruled out. Moreover, as actual neuronal connections between task-related regions of the STN, GPe, and SNr were not investigated in this current study, it would be desirable to investigate the actual connections between these areas by a method using gene manipulation with chemogenetics using the Designer Receptors Exclusively Activated by a Designer Drug (DREADD) method and examining changes in behavior.^48^

Second, we did not clarify the difference in function between the direct (from the striatum to SNr) and STN-GPe pathways, although both pathways are thought to be involved in facilitating behavior (Figure 5). Although the indirect and hyperdirect pathways are thought to contribute to action inhibition, the differences in their functions have not been clearly established. However, recent studies using genetic manipulation techniques in rodents have shown that the indirect pathway does not simply inhibit behavior as previously thought, but that it is involved in switching behavior or driving transient punishment.^49, 50^ Similarly, genetic manipulation and novel task-based behavioral experiments in rodents or non-human primates are needed to clarify the functional differences between the direct and STN-GPe pathways.

**Figure 5.**
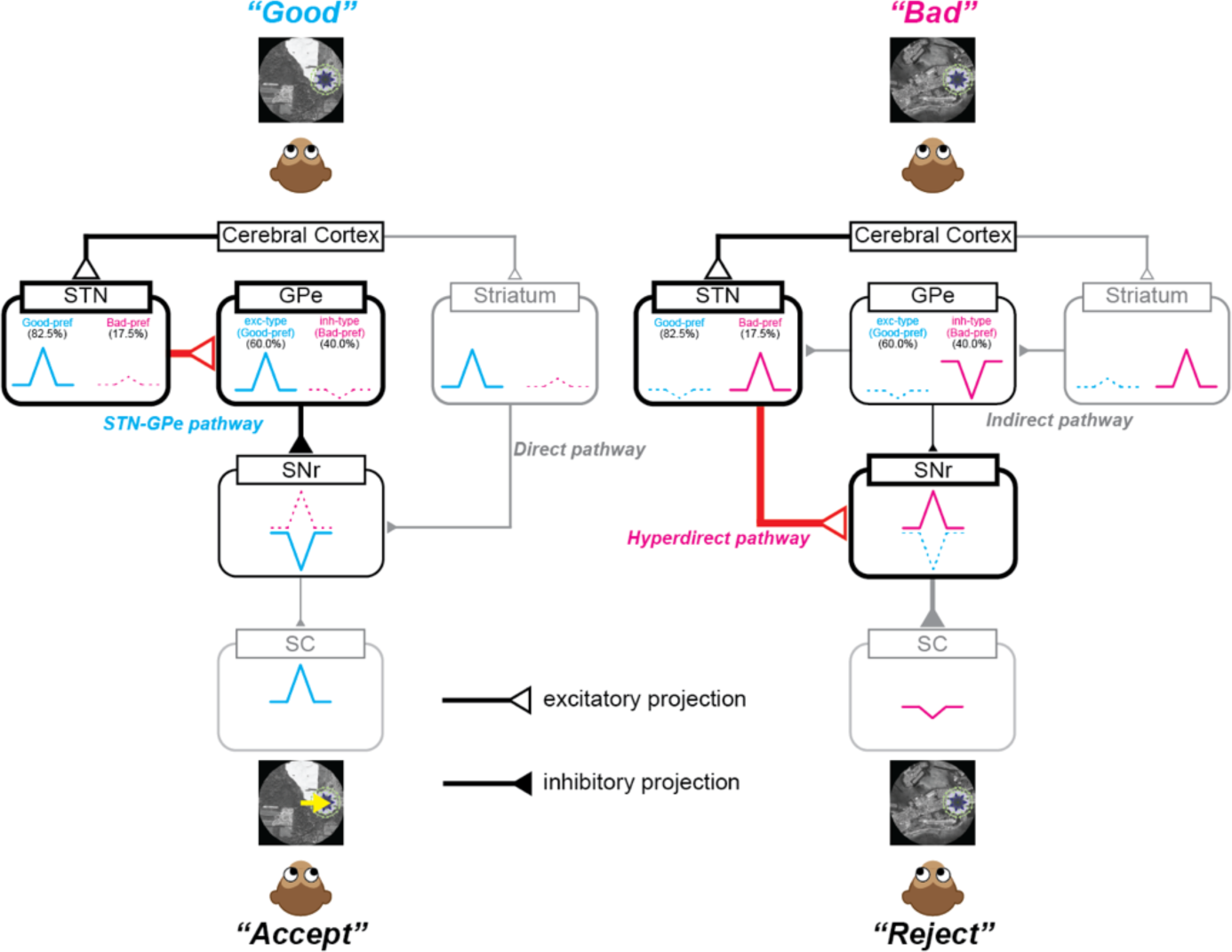
The new concept of the basal ganglia circuit. The new concept of the basal ganglia circuit based on results of this study. Abbreviations: GPe, globus pallidus external segment. SC, superior colliculus. SNpl, substantia nigra pars lateralis. SNr, substantia nigra pars reticulata. STN, subthalamic nucleus.

## Conclusions

In this study, we revealed that a subset of STN and GPe neurons significantly increased neural activity when the monkeys selected good objects. Local injection of a glutamatergic antagonist into the GPe resulted in delayed saccade reaction times toward contralateral good objects, whereas injection into the SNr accelerated saccade reaction times to contralateral bad objects. These findings suggest that excitatory projections from the STN to the GPe may play a role in promoting movement, while projections from the STN to the SNr may inhibit involuntary movements. As the STN-GPe circuit has not yet been accounted for in the three conventional basal ganglia pathways (direct, indirect, and hyperdirect), an additional pathway is likely to be involved.

## Methods

### Animal preparation

Two male macaque monkeys (*Macaca mulatta*, 9 years old, 9–10 kg body weight) referred to as C and S were used in this study. All animal care and experimental procedures were approved by the National Eye Institute Animal Care and Use Committee and complied with the Public Health Service Policy on Human Care and Use of Laboratory Animals. Following chair acclimation, a plastic head holder and recording chambers were implanted in the skull using ceramic screws and dental acrylic under general isoflurane anesthesia and aseptic conditions. Analgesics were administered intraoperatively and for several days thereafter. Oculomotor task training was initiated once the monkeys had fully recovered from the surgery. The daily water intake was controlled to motivate the monkeys to perform the behavioral tasks. In both the training and experimental sessions, the head of each monkey was fixed to a primate chair. Eye movements were monitored using an eye-tracking system (EyeLink 1000; SR Research, Ontario, Canada) at a sampling rate of 1000 Hz.

### Behavioral procedure

Behavioral tasks were controlled by a custom-made C++-based experimental data acquisition system (Blip; available at http://www.robilis.com/blip/). The monkeys sat on a primate chair facing a front screen in a dark sound-attenuated room. Visual stimuli generated by an active matrix liquid crystal display projector (PJ658, ViewSonic Corp., Brea, CA, USA) were projected onto the screen.

### Choice task

The present study used two tasks, choice and fixation tasks. In the choice task, large, circular (radius: 25°), gray scale aerial photographs derived from OpenAerialMap, referred to as *scenes,* were projected onto a screen in front of the subject as the background. Computer-generated multicolored fractal objects (radius: 5°) were created and used as the target.^51^ For each single-unit recording, one of six sets was randomly selected (Figure S1). Each set contained four *scenes,* with one good object (associated with reward delivery) and one bad object (associated with no reward delivery) in each *scene*. The object values remained unchanged in stable *scenes* 1 and 2, whereas their values changed in flexible *scenes* 3 and 4. This design allowed us to determine whether neuronal responses to objects were related to object features or their values.

The task procedure, illustrated in Figure 1A, began with the presentation of one of the four *scenes* after a trial interval of 1500–2000 ms. The subject was allowed to freely view the *scene* for 1000 ms; thereafter, a fixation point (FP) appeared at the center of the screen. If the subject maintained their gaze on the FP for 700 ms, a fractal object target was displayed randomly and sequentially after FP offset. In cases where the subject made a saccade toward the target and fixated on it for over 400 ms, it was interpreted as object acceptance, and a reward (0.4 mL) was delivered if the target was a good object or not delivered if it was a bad object. If the subject either continued gazing around the center point (*stay*), made a saccade to the target but returned their gaze to the center point within 400 ms (*return*), or looked away from the target (*other*), the FP reappeared after 1000 ms. The target range was set as a square with 8° per side. If the subject’s eye position moved outside the target range for <400 ms after making a saccade to the target, the target disappeared, and the FP was presented again. Consequently, when the monkeys rejected bad objects by *stay* or *other*, the waiting time for the next FP to appear was 1000 ms, whereas if they rejected bad objects by *return*, the waiting time was < 1000 ms. Upon gazing at the FP, another target was presented. This procedure was repeated up to 15 times until the subject selected a target object. The target was presented randomly at one of six positions (eccentricity: 15°; angles: 0°, 45°, 135°, 180°, 225°, and 315°).

### Fixation task

The objective of the fixation task was to investigate the involvement of each neuronal response in suppressing saccades toward the presented objects. After completion of the choice task, the fixation task was initiated if single neurons could be isolated. In this task, both fractal objects (one good and one bad object) used in *scene* 1 of the choice task were randomly and sequentially presented 2–4 times on either the left or right side, with each object displayed for 400 ms with an inter-object interval of 400 ms. The monkeys were required to maintain fixation on a white-colored FP while the objects were displayed on the screen simultaneously. A liquid reward (0.4 mL) was delivered 300 ms after the presentation of the final object. If the monkeys made a saccade toward any of the objects, the trial was deemed a fixation break error.

### MRI

Each monkey had two recording chambers implanted in its skull, one placed over the GPe and the other over the STN and SNr. Postoperatively, the monkeys underwent MRI to visualize the brain structures and grid holes using a gadolinium-based MRI contrast agent (Magnevist, Bayer Healthcare Pharmaceuticals, Wayne, NJ, USA), which was poured into the recording chambers. The recording sites were reconstructed based the MRI scans using a clinical 3T MR system (MAGNETOM Prisma; Siemens Healthcare, Erlangen, Germany). Both 3D T1-weighted (MPRAGE) and T2-weighted images (SPACE) with an isotropic voxel size of 0.5 mm were obtained.^52^ Additionally, QSM was performed to enhance the visualization of the STN, GPe, and SNr.^25, 53, 54^ QSM images were reconstructed from the phase images from the 3D multi-echo gradient-echo sequence with a repetition time of 50 ms and time to echo values of 3.7, 10.1, 16.7, 23.4, 30.0, 36.6, and 43.2 ms.^25^ The reconstruction process involved three main steps: phase image unwrapping, high-pass filtering of the phase image (background field removal), and dipole deconvolution (Figure 4a). These steps were executed using the morphology-enabled dipole inversion toolbox (http://pre.weill.cornell.edu/mri/pages/qsm.html)^55^ on MATLAB 2019 (The MathWorks, Inc., Natick, MA, USA).

### Visualization of STN, GPe, and SNr recording sites

To visualize the locations of the recorded task-related neurons in the STN, GPe, and SNr, QSM images were imported using 3D Slicer software (version 5.2.2).^56^ The regions of interest were manually created, converted into a 3D model, and aligned with the recording site using the Blender software (version 3.5). The number of recorded neurons was summarized every 500 μm in depth, and only the areas with three or more recorded neurons were noted (Figure S2).

### Neuronal recording procedure

Single-unit recordings were conducted using epoxy- or glass-coated tungsten electrodes with impedances ranging from 1–9 MΩ (Frederick Haer & Co., Bowdoinham, ME, USA; Alpha Omega Engineering, Nazareth, Israel). An oil-driven micromanipulator (MO-973A, Narishige, Tokyo, Japan) was used to introduce the electrode into the brain through a 23-gauge stainless-steel guide tube. Signals collected from the electrode were amplified and filtered (0.3–10 kHz; A-M Systems Inc., Carlsborg, WA, USA). The choice task was initiated once the neuronal activity was isolated. If the contralateral target onset modulated neuronal activity, the neuron was considered task-related, and recording continued. Two types of neurons were found in the STN and GPe: Good-pref and Bad-pref. These neuron types were manually classified by examining the peristimulus time histogram (PSTH) at the contralateral target onset of neuronal activity.

### Glutamatergic antagonist injection procedure

To inhibit excitatory projections from the STN into the GPe and SNr, we injected a mixture of CPP (C104, Sigma-Aldrich, Inc., St. Louis, MO, USA) and NBQX (N183, Sigma-Aldrich) into the caudal-dorsal part of the GPe or the lateral part of the SNr, where many task-related neurons were found. The glutamatergic antagonist was injected into either the right or left side of the GPe and the right side of the SNr of each monkey. Owing to the relatively high firing rate of the baseline activity in GPe and SNr neurons, it was possible to determine whether the tip of the electrode had reached the dorsal edge of the GPe and SNr. Prior to injection, neuronal activity was recorded using a custom-made injectrode.^57^

First, monkeys performed the choice and fixation tasks to collect pre-injection control data. Then, we injected 1 μL of 10–20 mM (for GPe) or 5–10 mM (for SNr) mixture of CPP and NBQX at 0.2 μL/min by a remotely controlled infusion pump (PHD ULTRA, Harvard Apparatus, Holliston, MA, USA). These concentrations were determined with reference to a previous study,^23^ while we used higher concentrations of CPP and NBQX (>5 mM) to investigate the effects of excitatory projections on behavior. After the injection was completed, the monkeys were required to repeatedly perform the choice and fixation tasks to assess the effect of the CPP and NBQX injections (from 5–90 min after injection). As a control, we injected 1 μL of saline into the same sites of the GPe and SNr where we injected the glutamatergic antagonists. For statistical analyses of the effect of the injection, we used data acquired 60–90 min after the injection of CPP, NBQX, and saline. However, only one of the data points from monkey S in the SNr (10 mM CPP and NBQX) was analyzed at 20 min post injection because the glutamatergic effect was too strong for the monkeys to continue performing the task.

### Data analysis

All behavioral and neurophysiological data were preprocessed using MATLAB 2020b (The MathWorks, Inc., Natick, MA, USA). For each neuron, data were aligned with the initiation of events (*scene*, target, and saccade onset). The neuronal response time course to event onset for each condition was examined by calculating PSTHs in 1-ms bins and smoothed with a spike density function using a Gaussian filter (σ = 20 ms). The activity in each neuron was Z-transformed to facilitate comparisons of neuronal activity across the three regions (STN, GPe, and SNr).^58, 59^ First, the baseline firing rate was calculated by averaging the firing rate during the 500-ms period before the *scene*, target, or saccade onset. This baseline was subtracted from the smoothed PSTH and aligned with the initiation of events. Subsequently, we performed a Z-score transformation for each PSTH by subtracting the mean baseline from the PSTH and dividing it by the SD of the PSTH.

The onset of saccades toward good and bad objects in the choice task was defined as the time when the eye speed exceeded 40°/s within 400 ms after target onset. In the fixation task, saccades toward objects were detected offline and fixation break errors were considered.

### Statistical analysis

No statistical methods were employed to predetermine the sample size. Instead, we consulted previous studies on the number of recorded neurons in the STN,^7^ GPe,^14^ and, SNr.^15^ Statistical analyses were conducted on the preprocessed data using R software (version 4.2.2),^60^ and the *lme4,*^61^ *pbkrtest,*^62^ and *emmeans*^63^ packages on RStudio.

We performed Welch’s t-test for each *scene* to compare saccade reaction times for good and bad objects (Figure 1E). We also conducted a Fisher’s exact test to determine whether the proportion of monkeys choosing *stay* for the bad object was larger in *scenes* 1 and 2, where the object values remained constant (Figure 1F).

To compare the normalized neuronal activity across the STN, GPe, and SNr, and the behavioral parameters such as saccade reaction times before and after injections, LMMs or generalized linear mixed-effects models (GLMMs) were used to prevent false positives.^64^ LMMs were employed for analyzing neuronal activity in the STN, GPe, and SNr under various conditions (Figures 2 and S3–S9) because the neural data were Z-transform normalized. Conversely, GLMMs were used to compare saccade reaction times and the proportion of chosen actions for bad objects in the choice task and fixation break errors in the fixation task (Figures 3 and S10) before and after injection.

For all statistical tests using the LMM and GLMM, we first compared the full model, containing explanatory variables as fixed effects. All monkey, neuron, or experimental session IDs were considered as random effects, which constituted the null model. We used a parametric bootstrap method to assess the models’ goodness of fit by performing 10,000 iterations and computing the *p*-value based on the difference in deviance between the two models. If the full model demonstrated a significant fit, we conducted post-hoc pairwise t-tests with Bonferroni correction to further explore these differences.

### 1) Comparison of normalized neuronal activity in STN, GPe, and SNr at contralateral object onset in the choice and fixation tasks (Figure 2)

We examined the differences in normalized neuronal activity for good and bad objects presented on the contralateral side within and across the choice and fixation tasks. The LMM was employed, and the full and null models used for comparison were as follows:

> *Full Model: NormalizedNeuronalActivity* ∼ *Value* × *Task* + (1|*monkey_ID*) + (1|*monkey_ID*: *Neuron_ID*),
>
> *Null Model: NormalizedNeuronalActivity* ∼ (1|*monkey_ID*) + (1|*monkey_ID*: *Neuron_ID*),

where *NormalizedNeuronalActivity* is the response variable, which is the mean Z-transformed PSTH of individual neurons during a 200 ms period, starting 100 ms after object onset in both the choice and fixation tasks, *Value* (good vs. bad), *Task* (choice vs. fixation), and their interaction are fixed effects, and *monkey_ID* (C and S) and *Neuron_ID* [Good-pref STN neurons, #1–81 (monkey C) and #1–70 (monkey S); Bad-pref STN neurons, #1–10 (monkey C) and #1– 22 (monkey S); Good-pref GPe neurons, #1–65 (monkey C) and #1–59 (monkey S); Bad-pref GPe neurons, #1–46 (monkey C) and #1–35 (monkey S); SNr neurons, #1–51 (monkey C) and #1–49 (monkey S)] are random effects to account for individual-level variability.

The level of statistical significance for the model comparison was set at α = 0.05. As we performed six pairwise t-tests for comparison among the normalized neuronal activity for good and bad objects in the choice and fixation tasks, the level of statistical significance for post-hoc tests was set at α = 0.05/6, applying a Bonferroni correction.

### 2) Comparison of normalized neuronal activity in the STN, GPe, and SNr at scene onset in the choice task (Figure S3)

We employed an LMM to evaluate the differences in normalized neuronal activity at *scene* onset. The full and null models used for the comparison were as follows:

> *Full Model: NormalizedNeuronalActivity* ∼ *Scene* + (1|*monkey_ID*) + (1|*monkey_ID*: *Neuron_ID*),
>
> *Null Model: NormalizedNeuronalActivity* ∼ (1|*monkey_ID*) + (1|*monkey_ID*: *Neuron_ID*),

where *NormalizedNeuronalActivity* is the mean Z-transformed PSTH of individual neurons during a 200 ms period, ranging from 100–300 ms after *scene* onset, *Scene* (1–4) is a fixed effect, and *monkey_ID* and *Neuron_ID* are random effects.

The level of statistical significance for the model comparison was set at α = 0.05. Post-hoc tests were not performed given that there was no significant difference among the *scenes* in each area.

### 3) Comparison of normalized neuronal activity in the STN, GPe, and SNr at target or saccade onset among four scenes in the choice task (Figures S4–S9)

We employed an LMM to investigate the differences in normalized neuronal activity for the target or saccade onset. The full and null models used for the comparison were as follows:

> *Full Model: NormalizedNeuronalActivity* ∼ *Scene* × *Value* × *Direction* + (1|*monkey_ID*) + (1|*monkey_ID*: *Neuron_ID*),
>
> *Null Model: NormalizedNeuronalActivity* ∼ (1|*monkey_ID*) + (1|*monkey_ID*: *Neuron_ID*),

where *NormalizedNeuronalActivity* is the mean Z-transformed PSTH of individual neurons during a 200 ms period, beginning 100 ms after target onset; *Scene* (1–4), *Value* (good vs. bad), and *Direction* (contralateral vs. ipsilateral) are fixed effects, and *monkey_ID* and *Neuron_ID* are random effects.

The level of statistical significance for the model comparison was set at α = 0.05. As we performed six pairwise t-tests for comparison of normalized neuronal activity for good and bad objects in the choice task within each *scene* and six pairwise t-tests for comparison across *scenes*, the level of statistical significance for the post-hoc test was set at α = 0.05/12, applying a Bonferroni correction.

### 4) Comparison of saccade reaction times in the choice task between before and after injection of glutamatergic antagonist in the GPe and SNr (Figures 3A)

Statistical analyses were conducted using a GLMM to compare saccade reaction times in the choice task before and after injection of glutamatergic antagonists into the GPe and SNr. Poisson distribution was employed to assess the before and after injection differences in saccade reaction times. In these analyses, the datasets from *scenes* 1–4 were combined and the median values were used as the response variable. The full and null models used for the comparison were as follows:

> *Full Model: MedianSaccadeReactionTimes* ∼ *Injection* × *PrePost* × *Value* × *Direction* + (1|*monkey_ID*) + (1|*monkey_ID*: *Session_ID*),
>
> *Null Model: MedianSaccadeReactionTimes* ∼ (1|*monkey_ID*) + (1|*monkey_ID*: *Session_ID*),

where *MedianSaccadeReactionTimes* is the median of saccade reaction times in each condition in the choice task, *Injection* (glutamatergic antagonist vs. saline), *PrePost* (pre-injection vs. post-injection), *Value* (good vs. bad), and *Direction* (contralateral vs. ipsilateral) are fixed effects, and *monkey_ID* and *Session_ID* [glutamatergic antagonist into GPe; #1–5 (monkey C) and #6–10 (monkey S); saline into GPe; #1–5 (monkey C) and #6–10 (monkey S); glutamatergic antagonist into SNr; #1–4 (monkey C) and #5–8 (monkey S); saline into SNr; #1–4 (monkey C) and #5–8 (monkey S)] are random effects.

Post-hoc tests were conducted before and after injection for each condition. The level of statistical significance for the model comparison was set at α = 0.05. As we performed eight pairwise t-tests to compare saccade reaction times, the level of statistical significance for the post-hoc test was set at α = 0.05/8, applying a Bonferroni correction.

### 5) Comparison of the proportions of chosen action for bad object in the choice task between before and after injection of glutamatergic antagonist in the GPe and SNr (Figures S10)

A GLMM with a binomial distribution was used for statistical analysis to compare the proportion of actions chosen for bad objects in the choice task before and after the injections. The full and null models used for the comparison were as follows:

> *Full Model: ChosenActionRate ((chosen action count) / (total trial count))* ∼ *Injection* × *PrePost* × *Value* × *Direction* + (1|*monkey_ID*) + (1|*monkey_ID*: *Session_ID*), weights = (total trial count),
>
> *Null Model: ChosenActionRate ((chosen action count) / (total trial count))* ∼ (1|*monkey_ID*) + (1|*monkey_ID*: *Session_ID*), weights = (total trial count)

where *ChosenActionRate* is the proportion of chosen actions for bad objects in each condition in the choice task; *Injection* (glutamatergic antagonist vs. saline), *PrePost* (pre-injection vs. post-injection), *Value* (good vs. bad), and *Direction* (contralateral vs. ipsilateral) are fixed effects; and *monkey_ID* and *Session_ID* are random effects for model comparison.

Post-hoc tests were conducted before and after injection for each condition. The level of statistical significance for the model comparison was set at α = 0.05. As we performed eight pairwise t-tests to compare the proportions of chosen actions for bad objects, the level of statistical significance for the post-hoc test was set at α = 0.05/8, applying a Bonferroni correction.

### 6) Comparison of the proportion of fixation break errors in the fixation task between before and after injection of glutamatergic antagonist in the GPe and SNr (Figures 3B)

A GLMM with binominal distribution was used for statistical analysis to compare the proportion of fixation break errors in the fixation task before and after the injections. The full and null models used for comparison were as follows:

> *Full Model: FixBreakErrorRate ((error trial count) / (total trial count))* ∼ *Injection* × *PrePost* × *Value* × *Direction* + (1|*monkey_ID*) + (1|*monkey_ID*: *Session_ID*), weights = (total trial count),
>
> *Null Model: FixBreakErrorRate ((error trial count) / (total trial count))* ∼ (1|*monkey_ID*) + (1|*monkey_ID*: *Session_ID*), weights = (total trial count)

where *FixBreakErrorRate* is the proportion of fixation break errors in each condition in the fixation task, *Injection* (glutamatergic antagonist vs. saline), *PrePost* (pre-injection vs. post-injection), *Value* (good vs. bad), and *Direction* (contralateral vs. ipsilateral) are fixed effects, and *monkey_ID* and *Session_ID* are random effects.

Post-hoc tests were conducted before and after injection for each condition. The level of statistical significance for the model comparison was set at α = 0.05. As we performed eight pairwise t-tests to compare the proportions of chosen actions for bad objects, the level of statistical significance for the post-hoc test was set at α = 0.05/8, applying a Bonferroni correction.

## Acknowledgments

This research was supported by the Intramural Research Program at the National Institutes of Health, National Eye Institute (1ZIA EY000415). MRI scanning was conducted in the Neurophysiology Imaging Facility Core (National Institute of Mental Health, National Institute of Neurological Disorders and Stroke, and National Eye Institute). We thank D. Parker, H. Warnock, G. Tansey, K. Allen-Worthington, A. MacLarty, M.K. Smith, A.M. Nichols, D. Yochelson, A.V. Hays, J. Fuller-Deets, and M. Robinson for technical assistance.

## Author contributions

Conceptualization, A.Y.; Software, A.Y.; Formal Analysis, A.Y.; Writing – Original Draft, A.Y.; Writing – Review & Editing, A.Y.; Visualization, A.Y.; Supervision O.H.; Funding Acquisition, O.H.

## Resource availability

Further information and requests for resources and reagents should be directed to and will be fulfilled by the lead contact, Atsushi Yoshida.

## Materials availability

This study did not generate new unique reagents.

## Data and code availability

Data and code are available upon request.

## Supplemental information

Figures S1–S10, Tables S1–S15

## References

1. Albin, R.L., Young, A.B., and Penney, J.B. (1989). The functional anatomy of basal ganglia disorders. Trends Neurosci. 12, 366–375.

2. DeLong, M.R. (1990). Primate models of movement disorders of basal ganglia origin. Trends Neurosci. 13, 281–285.

3. Hikosaka, O., Takikawa, Y., and Kawagoe, R. (2000). Role of the basal ganglia in the control of purposive saccadic eye movements. Physiol. Rev. 80, 953–978.

4. Mink, J.W. (2003). The Basal Ganglia and involuntary movements: impaired inhibition of competing motor patterns. Arch. Neurol. 60, 1365–1368.

5. Nambu, A. (2008). Seven problems on the basal ganglia. Curr. Opin. Neurobiol. 18, 595– 604.

6. Nambu, A., Tokuno, H., and Takada, M. (2002). Functional significance of the cortico-subthalamo-pallidal “hyperdirect” pathway. Neurosci. Res. 43, 111–117.

7. Pasquereau, B., and Turner, R.S. (2017). A selective role for ventromedial subthalamic nucleus in inhibitory control. Elife. 6, e31627

8. Mosher, C.P., Mamelak, A.N., Malekmohammadi, M., Pouratian, N., and Rutishauser, U. (2021). Distinct roles of dorsal and ventral subthalamic neurons in action selection and cancellation. Neuron. 109, 869–881.

9. London, D., Fazl, A., Katlowitz, K., Soula, M., Pourfar, M.H., Mogilner, A.Y., and Kiani, R. (2021). Distinct population code for movement kinematics and changes of ongoing movements in human subthalamic nucleus. Elife. 10, e64893.

10. Brotchie, P., Iansek, R., and Horne, M.K. (1991). Motor function of the monkey globus pallidus. 1. Neuronal discharge and parameters of movement. Brain. 114, 1667–1683.

11. Mitchell, S.J., Richardson, R.T., Baker, F.H., and DeLong, M.R. (1987). The primate globus pallidus: neuronal activity related to direction of movement. Exp. Brain Res. 68, 491–505.

12. Turner, R.S., and Anderson, M.E. (1997). Pallidal discharge related to the kinematics of reaching movements in two dimensions. J. Neurophysiol. 77, 1051–74.

13. Yoshida, A., and Tanaka, M. (2009). Enhanced modulation of neuronal activity during antisaccades in the primate globus pallidus. Cereb. Cortex. 19, 206–217.

14. Yoshida, A., and Tanaka, M. (2016). Two Types of Neurons in the Primate Globus Pallidus External Segment Play Distinct Roles in Antisaccade Generation. Cereb. Cortex. 26, 1187–1199.

15. Shin, S., and Sommer, M.A. (2010). Activity of neurons in monkey globus pallidus during oculomotor behavior compared with that in substantia nigra pars reticulata. J. Neurophysiol. 103,1874–1887.

16. Goldberg, J.A., and Bergman, H. (2011). Computational physiology of the neural networks of the primate globus pallidus: function and dysfunction. Neuroscience. 198, 171–192.

17. van der Kooy, D., Hattori, T., Shannak, K., and Hornykiewicz, O. (1981). The pallido-subthalamic projection in rat: anatomical and biochemical studies. Brain Res. 204, 253–268.

18. Moriizumi, T., and Hattori, T. (1992). Neuroscience. Separate neuronal populations of the rat globus pallidus projecting to the subthalamic nucleus, auditory cortex and pedunculopontine tegmental area. Neuroscience. 46, 701–710.

19. Shink, E., Bevan, M.D., Bolam, J.P., and Smith, Y. (1996). The subthalamic nucleus and the external pallidum: two tightly interconnected structures that control the output of the basal ganglia in the monkey. Neuroscience. 73, 335–357.

20. Nambu, A., Tokuno, M., Hamada, I., Kita, H., Imanishi, M., Akazawa, T., Ikeuchi, Y., and Hasegawa, N. (2000). Excitatory cortical inputs to pallidal neurons via the subthalamic nucleus in the monkey. J. Neurophysiol. 84, 289–300.

21. Kita, H., Nambu, A., Kaneda, K., Tachibana, Y., and Takada, M. (2004). Role of ionotropic glutamatergic and GABAergic inputs on the firing activity of neurons in the external pallidum in awake monkeys. J. Neurophysiol. 92, 3069–3084.

22. Hikosaka, O., and Wurtz, R.H. (1985). Modification of saccadic eye movements by GABA-related substances. Ⅰ. Effect of muscimol and bicuculline in monkey superior colliculus. J. Neurophysiol. 53, 266–291.

23. Tachibana, Y., Iwamuro, H., Kita, H., Takada, M., and Nambu, A. (2011). Subthalamo-pallidal interactions underlying parkinsonian neuronal oscillations in the primate basal ganglia. Eur. J. Neurosci. 34, 1470–1484.

24. Allen, M., Poggiali, D., Whitaker, K., Marshall, T.R., van Langen, J., and Kievit, R.A. (2021). Raincloud plots: a multi-platform tool for robust data visualization. Wellcome Open Research. 4, 63.

25. Yoshida, A., Ye, F.Q., Yu, D.K., Leopold, D.A., and Hikosaka, O. (2012). Visualization of iron-rich subcortical structures in non-human primates in vivo by quantitative susceptibility mapping at 3T MRI. Neuroimage. 241, e118429.

26. Francois, C., Percheron, G., and Yelnik, J. (1984). Localization of nigrostriatal, nigrothalamic and nigrotectal neurons in ventricular coordinates in macaques. Neuroscience. 13, 61–76.

27. Francois, C., Percheron, G., Yelnik, J., and Heyner, S. (1985). A histological atlas of the macaque (Macaca mulatta) substantia nigra in ventricular coordinates. Brain Res. Bull. 14, 349–367

28. Ma, T. P. (1989). Identification of the substantia nigra pars lateralis in the macaque using cytochrome oxidase and fiber stains. Brain Res. 480, 305–311.

29. Saleem, K.S., Avram, A.V., Glen, D., Yen, C.C., Ye, F.Q., Komlosh, M., and Basser, P.J. (2021). High-resolution mapping and digital atlas of subcortical regions in the macaque monkey based on matched MAP-MRI and histology. Neuroimage. 245, e118759.

30. Matsumura, M., Kojima, J., Gardiner, T.W., and Hikosaka, O. (1992) Visual and oculomotor functions of monkey subthalamic nucleus. J. Neurophysiol. 67, 1615–1632.

31. Isoda, M., and Hikosaka, O. (2008). Role for subthalamic nucleus neurons in switching from automatic to controlled eye movement. J. Neurosci. 28, 7209–7218.

32. Stanton, G.B., Goldberg, M.E., and Bruce, C.J. (1988). Frontal eye field efferents in the macaque monkey: I. Subcortical pathways and topography of striatal and thalamic terminal fields. J. Comp. Neurol. 271, 473–492.

33. Shook, B.L., Schlag-Rey, M., and Schlag, J. (1990). Primate supplementary eye field: I. Comparative aspects of mesencephalic and pontine connections. J. Comp. Neurol. 301, 618–642.

34. Kim, H.F., Amita, H., and Hikosaka, O. (2017). Indirect Pathway of Caudal Basal Ganglia for Rejection of Valueless Visual Objects. Neuron. 94, 920–930.

35. Parent, A., and Smith, Y. (1987). Organization of efferent projections of the subthalamic nucleus in the squirrel monkey as revealed by retrograde labeling methods. Brain Res. 436, 296–310.

36. Deniau, J.M., Hammond, C., Chevalier, G., and Feger, J. (1978). Evidence for branched subthalamic nucleus projections to substantia nigra, entopeduncular nucleus and globus pallidus. Neurosci. Lett. 9, 117–121.

37. Temel, Y., Visser-Vandewalle, V., and Carpenter, R.H.S. (2008). Saccadic latency during electrical stimulation of the human subthalamic nucleus. Curr. Biol. 18, 412–414.

38. Yugeta, A., Terao, Y., Fukuda, H., Hikosaka, O., Yokochi, F., Okiyama, R., Taniguchi, M., Takahashi, H., Hamada, I., Hanajima, R., and Ugawa, Y. (2010). Effects of STN stimulation on the initiation and inhibition of saccade in Parkinson disease. Neurology. 74, 743–748.

39. van den Wildenberg, W.P.M., van Wouwe, N.C., Ridderinkhof, K.R., Neimat, J.S., Elias, W.J., Bashore, T.R., and Wylie, S.A. (2021). Deep-brain stimulation of the subthalamic nucleus improves overriding motor actions in Parkinson’s disease. Behav. Brain Res. 402, 113124.

40. Hikosaka, O., and Wurtz, R.H. (1983). Visual and oculomotor functions of monkey substantia nigra pars reticulata. Ⅰ. Relation of visual and auditory responses to saccades. J. Neurophysiol. 49, 1230–1253.

41. Handel, A., and Glimcher, P.W. (1999). Quantitative analysis of substantia nigra pars reticulata activity during a visually guided saccade task. J. Neurophysiol. 82, 3458–3475.

42. Sato, M., and Hikosaka, O. (2002). Role of primate substantia nigra pars reticulata in reward-oriented saccadic eye movement. J. Neurosci. 22, 2363–2373.

43. Yelnik, J., Francois, C., Percheron, G., and Heyner, S. (1987). Golgi study of the primate substantia nigra. Ⅰ. Quantitative morphology and typology of nigral neurons. J. Comp. Neurol. 265, 455–472.

44. Francois, C., Yelinik, J., and Percheron, G. (1987). Golgi study of the primate substantia nigra. Ⅱ. Spatial organization of dendritic arborizations in relation to the cytoarchitectonic boundaries and to the striatonigral bundle. J. Comp. Neurol. 265, 473–493.

45. Beckstead, R.M., Edwards, S.B., and Frankfurter, A. (1981). A comparison of the intranigral distribution of nigratectal neurons labeled with horseradish peroxidase in the monkey, cat, ad rat. J. Neurosci. 2, 121–125.

46. May, P.J., and Hall, W.C. (1986). The sources of the nigrotectal pathway. Neuroscience. 19, 159–180.

47. Parent, A., and Bellefeullille, L.D. (1983). The pallidointralaminar and pallidonigra projections in primate as studied by retrograde double-labeling method. Brain Res. 278, 11–27.

48. Hasegawa, T., Chiken, S., Kobayashi, K., and Nambu, A. (2022). Subthalamic nucleus stabilizes movements by reducing neural spike variability in monkey basal ganglia. Nat. Commun. 13, 2233.

49. Nonomura, S., Nishizawa, K., Sakai, Y., Kawaguchi, Y., Kato, S., Uchigashima, M., Watanabe, M., Yamanaka, K., Enomoto, K., Chiken, S, Sano, H., Soma, S., Yoshida, J., Semejima, K., Ogawa, M., Kobayashi, K., Nambu, A., Isomura, Y., and Kimura, M. (2018). Monitoring and Updating of Action Selection for Goal-Directed Behavior through the Striatal Direct and Indirect Pathways. Neuron. 99, 1302–1314.

50. Isett, B.R., Nguyen, K.P., Schwenk, J.C., Yurek, J.R., Snyder, C.N., Vounatsos, M.V., Adegbesan, K.A., Ziausyte, U., and Gittis, A.H. (2023). The indirect pathway of the basal ganglia promotes transient punishment but not motor suppression. Neuron. S0896–6273.

51. Yamamoto, S., Kim, H.F., and Hikosaka, O. (2013). Reward value-contingent changes of visual responses in the primate caudate tail associated with a visuomotor skill. J. Neurosci. 33, 11227–11238.

52. Mugler, J.P. 3rd, Bao, S., Mulkern, R.V., Guttmann, C.R., Robertson, R.L., Jolesz, F.A., and Brookeman, J.R. (2000). Optimized single-slab three-dimensional spin-echo MR imaging of the brain. Radiology. 216, 891–899.

53. Liu, C., Li, W., Tong, K.A., Yeom, K.W., and Kuzminski, S. (2015). Susceptibility-weighted imaging and quantitative susceptibility mapping in the brain. J. Magn. Reson. Imaging. 42, 23–41.

54. Wang, Y., and Liu, T. (2015). Quantitative susceptibility mapping (QSM): decoding MRI data for a tissue magnetic biomarker. Magn. Reson. Med. 73, 82–101.

55. Liu, J., Liu, T., de Rochefort, L., Ledoux, J., Khalidov, I., Chen, W., Tsiouris, A.J., Wisnieff, C., Spincemaille, P., Prince, M.R., and Wang, Y. (2012). Morphology enabled dipole inversion for quantitative susceptibility mapping using structural consistency between the magnitude image and the susceptibility map. Neuroimage. 59, 2560–2568.

56. Fodorov, A., Beichel, R., Kalpathy-Cramer, J., Finet, J., Fillion-Robin, L.C., Pujol, S., Bauer, C., Jennings, D., Fennessy, F.M., Sonka, M., Buatti, J., Aylward, S.R., Miller, J.V., Pieper, S., and Kikinis, R. (2012). 3D Slicer as an Image Computing Platform for the Quantitative Imaging Network. Magn. Reson. Imaging. 30, 1323–1341.

57. Vanegas, M.I., Hubbard, K.R., Esfandyarpour, R., and Noudoost, B. (2019). Microinjectrode System for Combined Drug Infusion and Electrophysiology. J. Vis. Exp. 153, e60365.

58. Adler, A., Katabi, S., Finkes, I., Israel, Z., Prut, Y., and Bergman, H. (2012). Temporal convergence of dynamic cell assemblies in the striato-pallidal network. J. Neurosci. 32, 2473–2484.

59. Kaplan, A., Mizrahi-Kliger, A.D., Israel, Z., Adler, A., and Bergman, H. (2020). Dissociable roles of ventral pallidum neurons in the basal ganglia reinforcement learning network. Nat. Neurosci. 23, 556–564.

60. R Development Core Team (2020). R: A language and environment for statistical computing (Vienna, Austria: R Foundation for Statistical Computing).

61. Bates, D., Ma€chler, M., Bolker, B., and Walker, S. (2014). Fitting linear mixed-effects models using lme4. J. Stat. Softw. 67, 1–48.

62. Halekoh, U., and Højsgaard, S. (2014). A kenward-roger approximation and parametric bootstrap methods for tests in linear mixed models–the R package pbkrtest. J. Stat. Softw. 59, 1–30.

63. Lenth, R., Singmann, H., Love, J., Buerkner, P., and Herve, M. (2019). Estimated marginal means, aka least-squares means. R package version 1.3.2. https://rdrr.io/cran/emmeans/.

64. Yu, Z., Guindani, M., Grieco, S.F., Chen, L., Holmes, T.C., Xu, X. (2022). Beyond t test and ANOVA: applications of mixed-effects models for more rigorous statistical analysis in neuroscience research. Neuron. 110, 21–35.

