## Supplemental Figures & Tables for "Opposing functions of glutamatergic inputs between the globus pallidus external segment and substantia nigra pars reticulata"

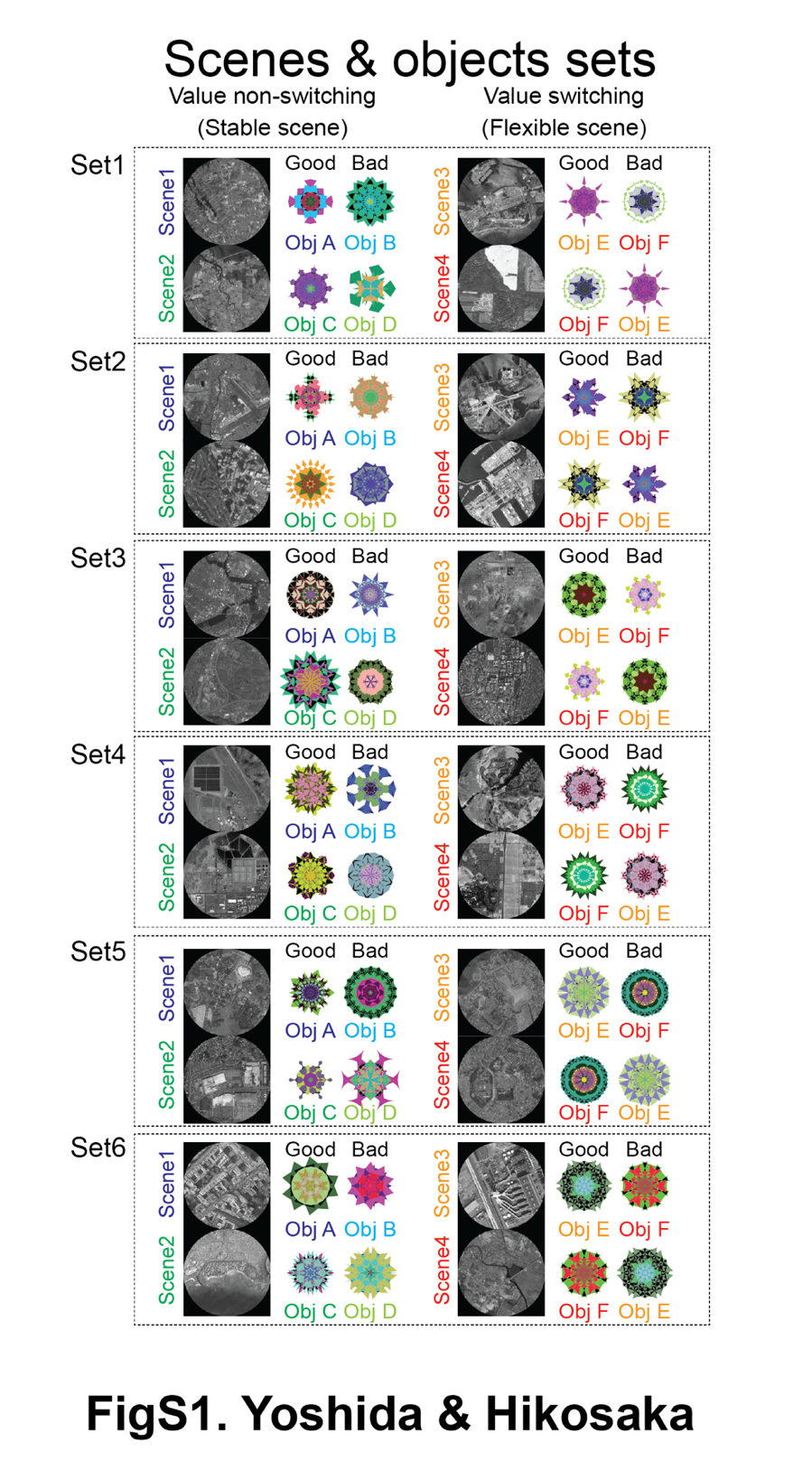
***Supplementary Figures***

**Figure S1 All sets of scenes 1-4 and good and bad objects for the choice task**

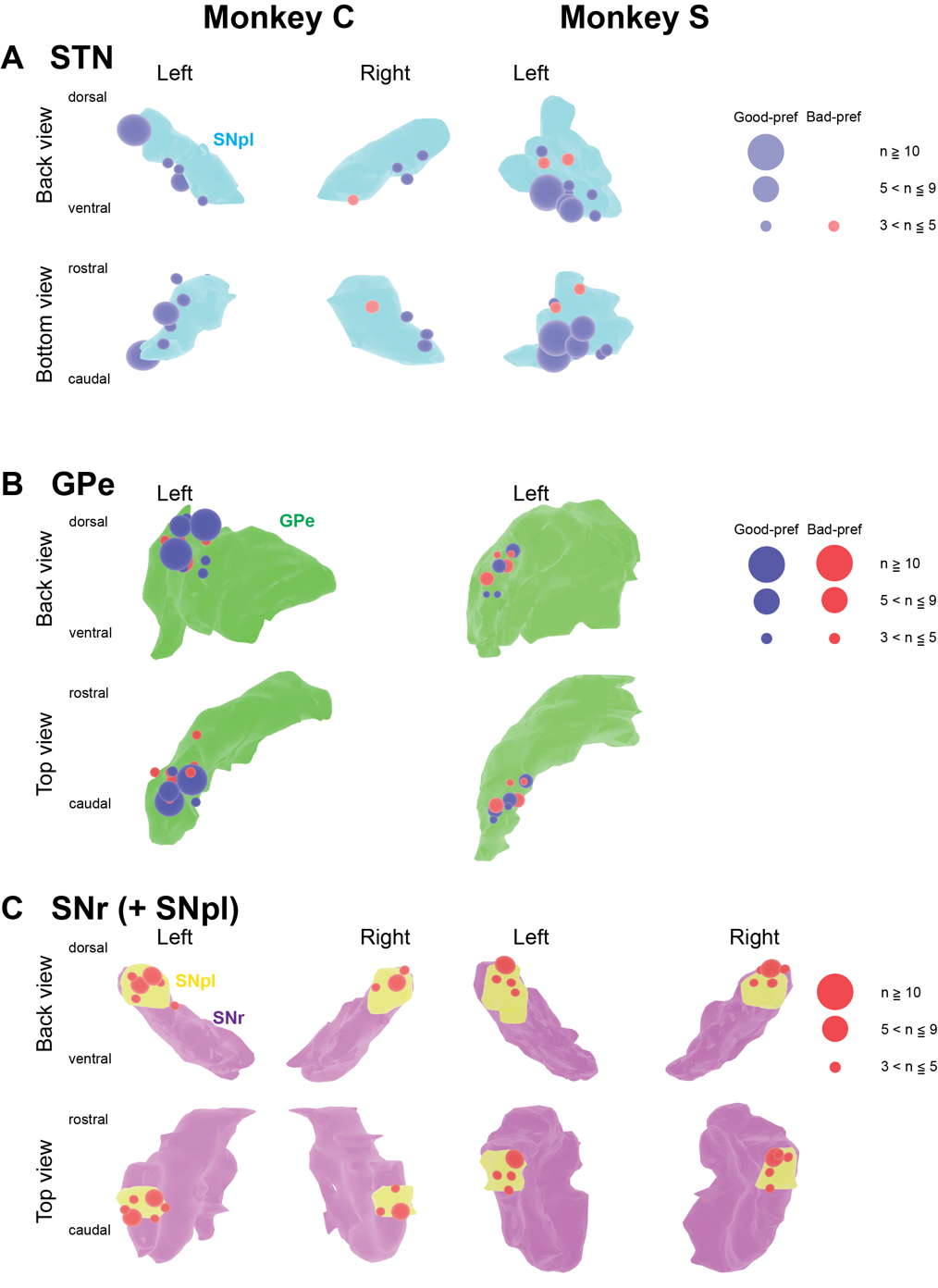
**Figure S2 Recording sites of task-related neurons in the STN, GPe, and SNr of two monkeys.**

Abbreviations: GPe, globus pallidus external segment. SNpl, substantia nigra pars lateralis. SNr, substantia nigra pars reticulata. STN, subthalamic nucleus.

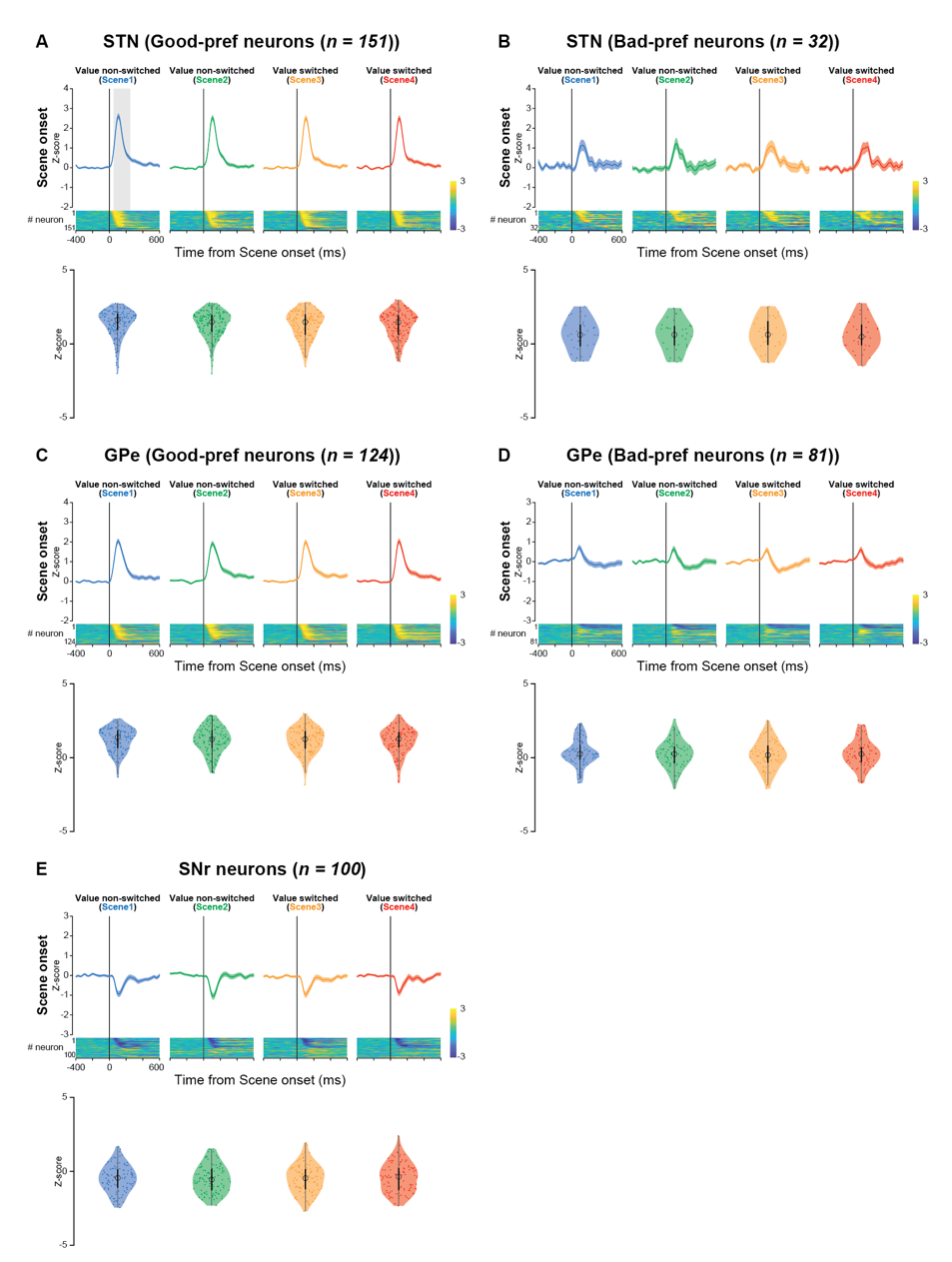
 **Figure S3 Population activity of Good-pref and Bad-pref STN neurons, Good-pref and Bad-pref GPe neurons, and SNr neurons at scene onset in the choice task**

Normalized mean population activity of neurons aligned at *scene* onset and the color maps of the normalized firing rate of individual neurons during the choice task. The violone plots of the mean normalized firing rate of individual neurons. Neuronal activity was measured for a 200-ms interval from 50 ms after *scene* onset (gray rectangle in (A)).

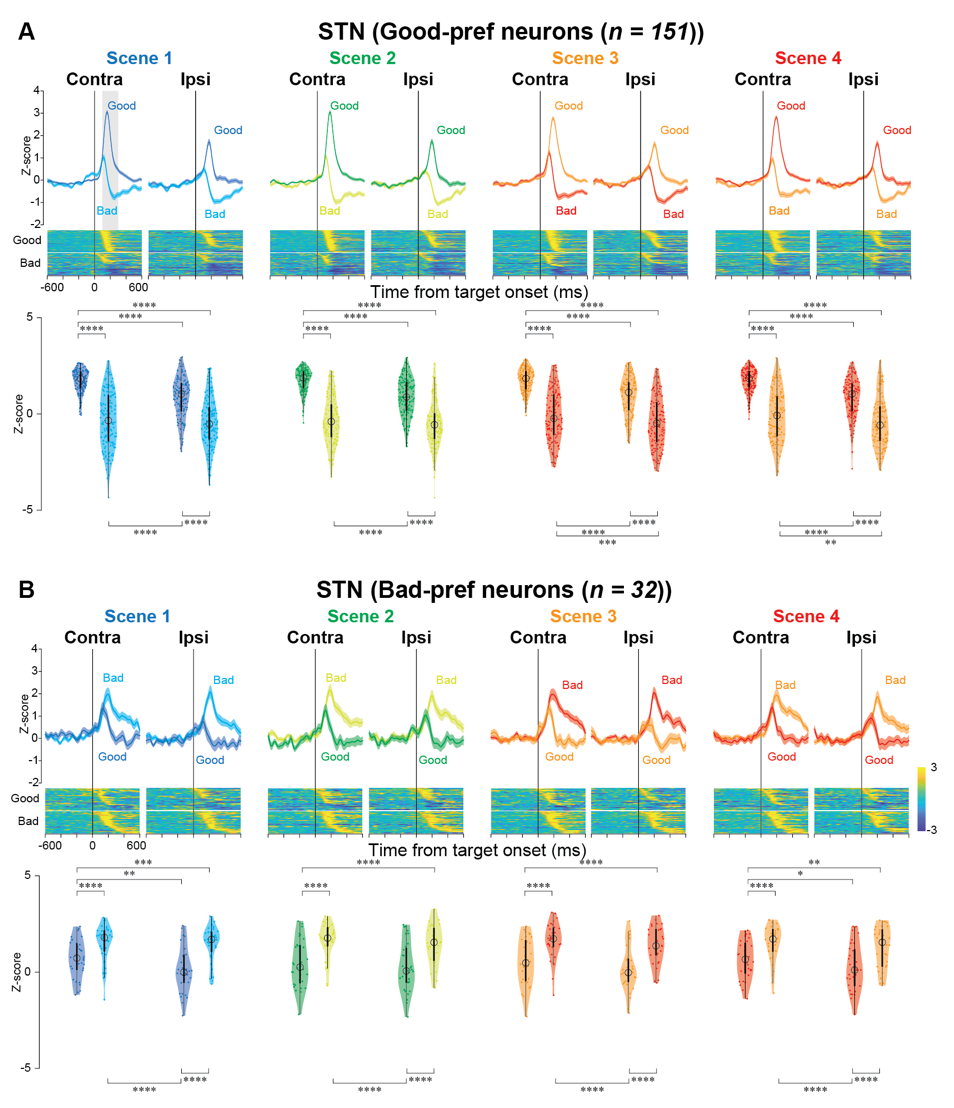
 **Figure S4 Population activity of Good-pref and Bad-pref STN neurons at contralateral and ipsilateral target onset in the choice task**

(A, B) Normalized mean population activity of Good- and Bad-pref STN neurons aligned at contralateral and ipsilateral target onset and the color maps of the normalized firing rate of individual neurons during the choice task. The violone plots of the mean normalized firing rate of individual neurons. Neuronal activity was measured for a 200-ms interval from 100 ms after target onset (gray rectangle in (A)). The asterisk indicates a significant difference in normalized neuronal activity in comparison among the conditions in the choice tasks (post hoc pairwise t-tests with Bonferroni correction, **p <* 0.05, ***p* < 0.01, ****p* < 0.001, *****p* < 0.0001). Asterisks are attached only to combinations with significant differences.

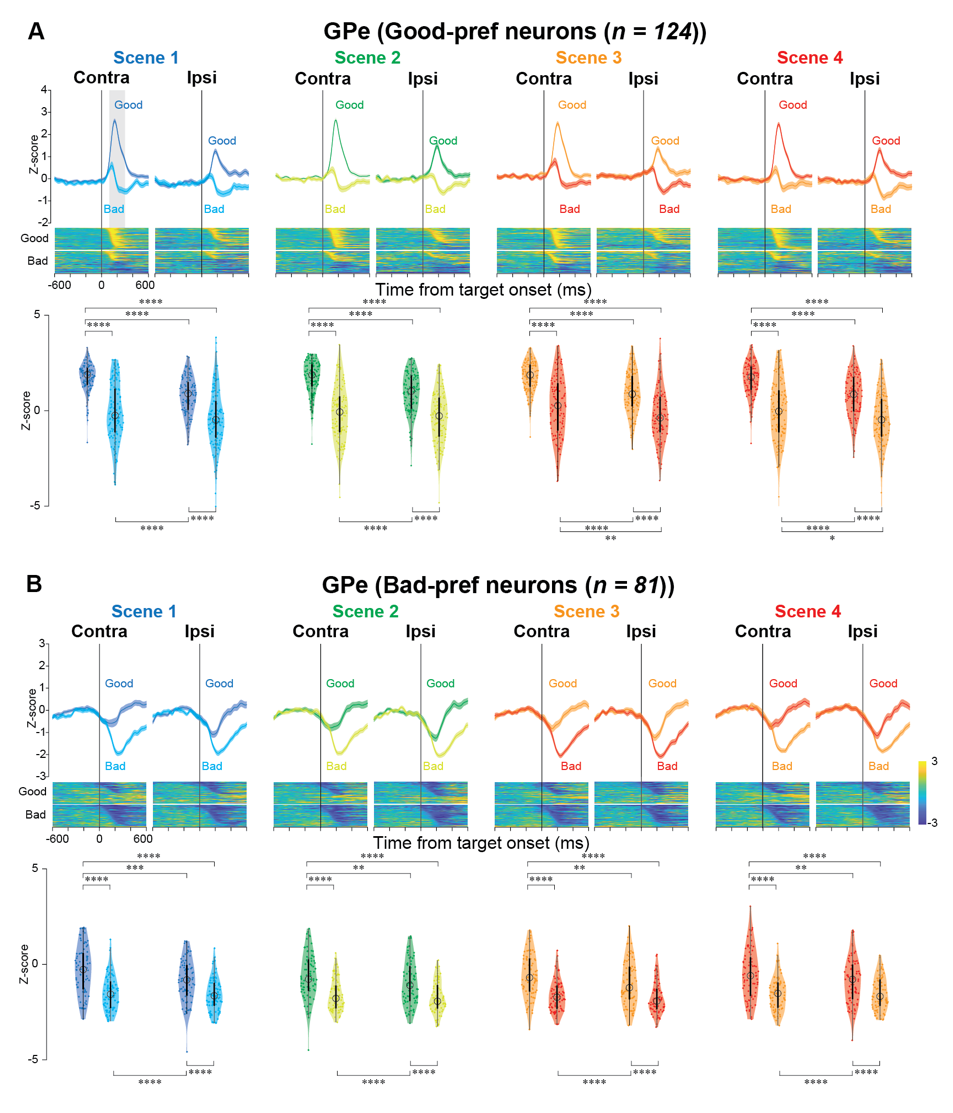
**Figure S5 Population activity of Good-pref and Bad-pref GPe neurons at contralateral and ipsilateral target onset in the choice task**

(A, B) Normalized mean population activity of Good- and Bad-pref GPe neurons aligned at contralateral and ipsilateral target onset and the color maps of the normalized firing rate of individual neurons during the choice task. The descriptions of this figure are the same as in Figure S3.

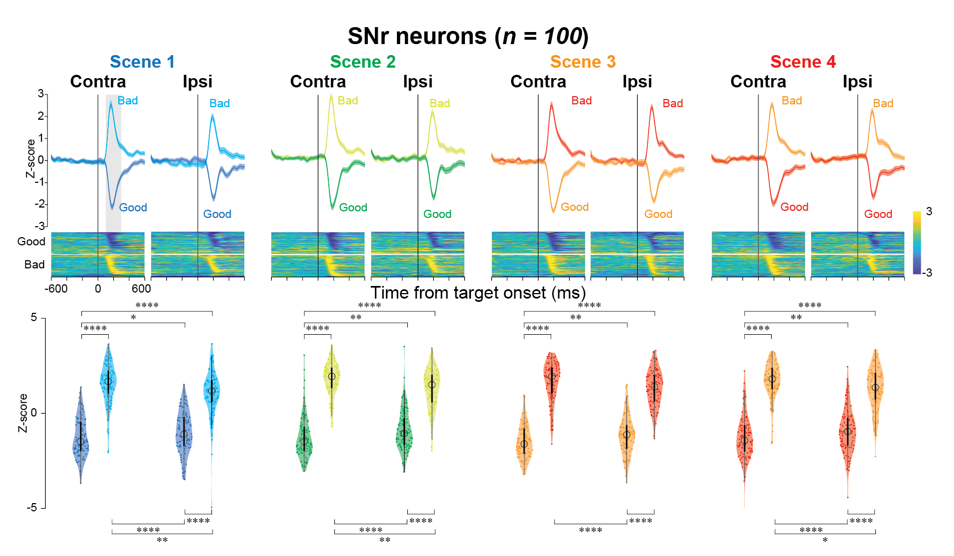
**Figure S6 Population activity of SNr neurons at contralateral and ipsilateral target onset in the choice task**

Normalized mean population activity of SNr neurons aligned at contralateral and ipsilateral target onset and the color maps of the normalized firing rate of individual neurons during the choice task. The descriptions of this figure are the same as in Figure S3.

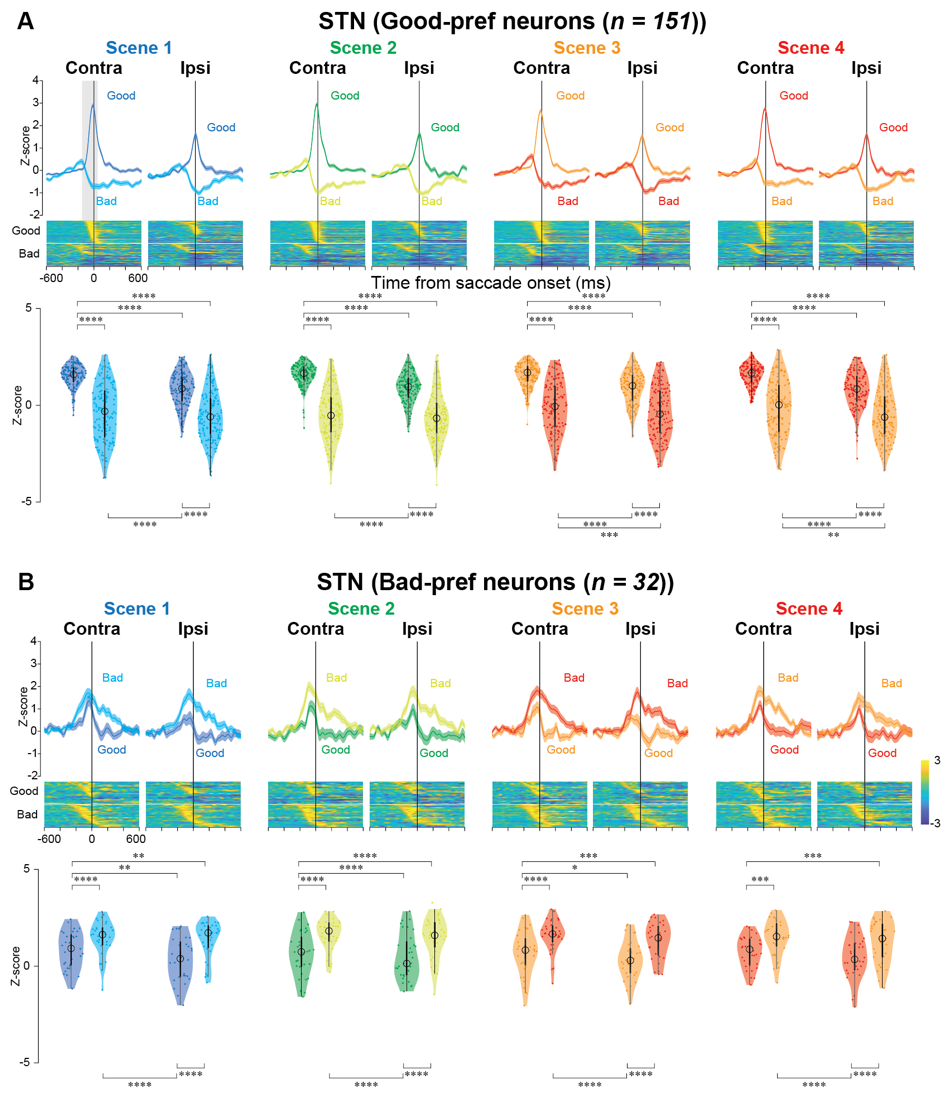
 **Figure S7 Population activity of Good-pref and Bad-pref STN neurons at contralateral and ipsilateral saccade onset in the choice task**

(A, B) Normalized mean population activity of Good- and Bad-pref STN neurons aligned at contralateral and ipsilateral saccade onset and the color maps of the normalized firing rate of individual neurons during the choice task. The violone plots of the mean normalized firing rate of individual neurons. Neuronal activity was measured for a 200-ms interval from 100 ms after saccade onset (gray rectangle in (A)). The asterisk indicates a significant difference in normalized neuronal activity in comparison among the conditions in the choice tasks (post hoc pairwise t-tests with Bonferroni correction, **p <* 0.05, ***p* < 0.01, ****p* < 0.001, *****p* < 0.0001). Asterisks are attached only to combinations with significant differences.

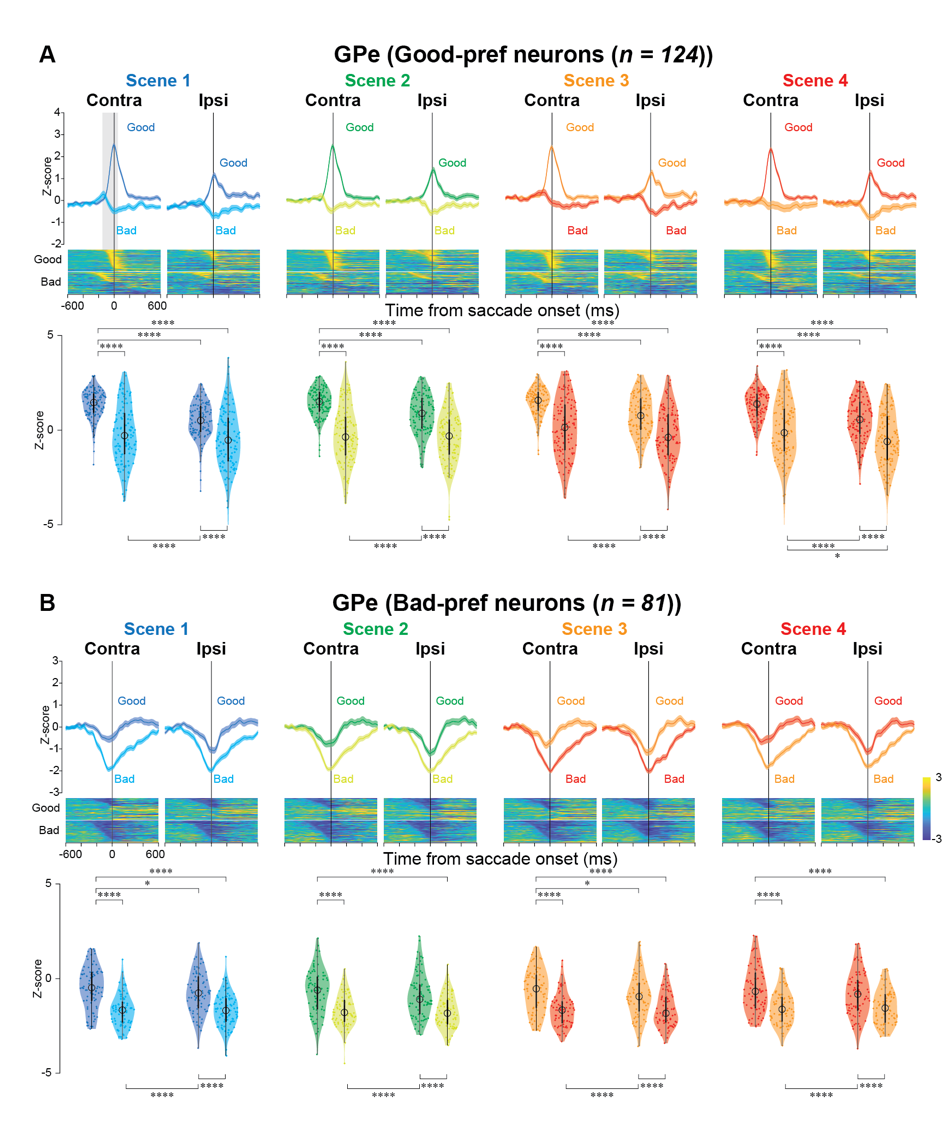
**Figure S8 Population activity of Good-pref and Bad-pref GPe neurons at contralateral and ipsilateral saccade onset in the choice task**

(A, B) Normalized mean population activity of Good- and Bad-pref GPe neurons aligned at contralateral and ipsilateral saccade onset and the color maps of the normalized firing rate of individual neurons during the choice task. The descriptions of this figure are the same as in Figure S6.

­­

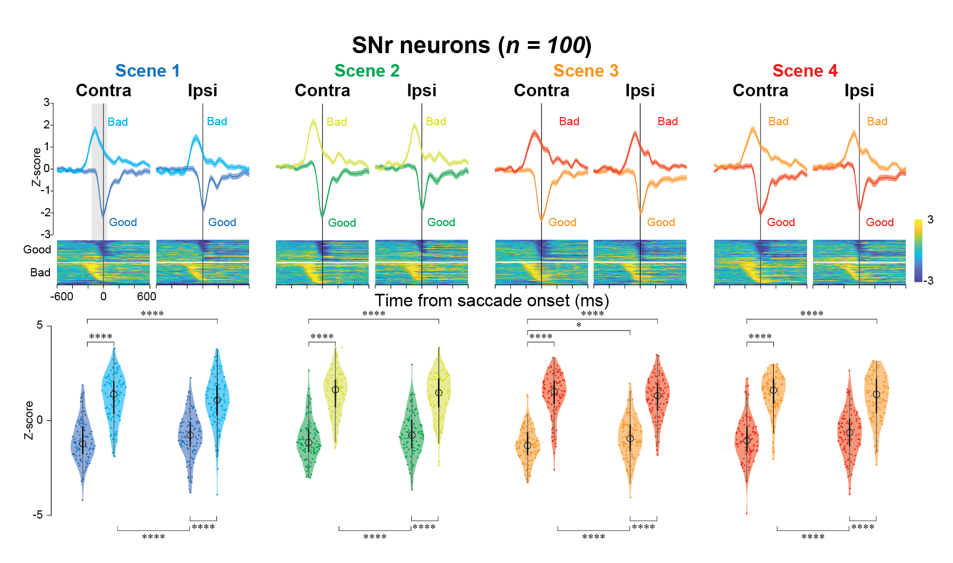
**Figure S9 Population activity of SNr neurons at contralateral and ipsilateral saccade onset in the choice task**

Normalized mean population activity of SNr neurons aligned at contralateral and ipsilateral saccade onset and the color maps of the normalized firing rate of individual neurons during the choice task. The descriptions of this figure are the same as in Figure S6.

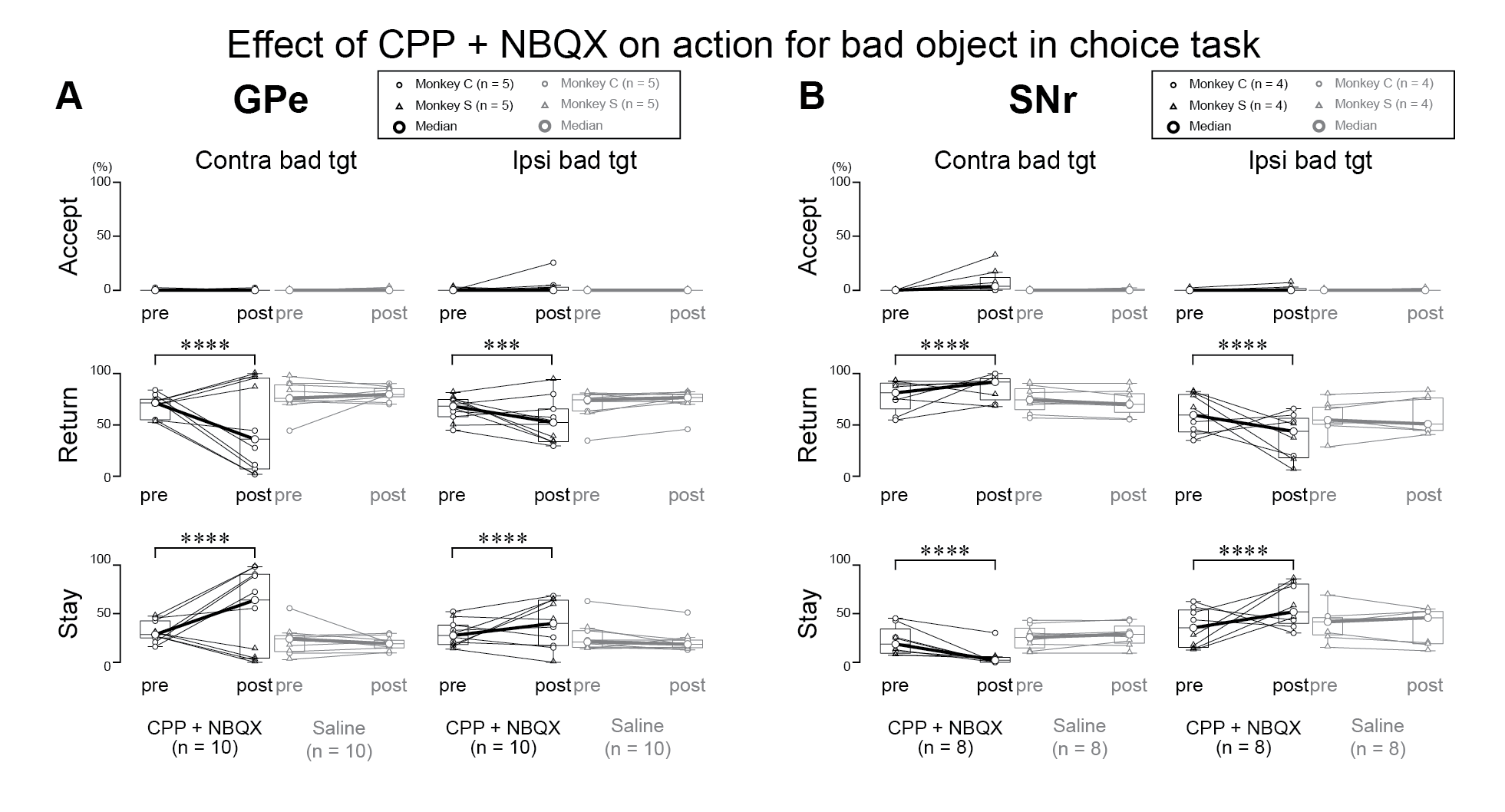
**Figure S10 Effects of glutamatergic antagonist injection into GPe and SNr on the proportion of chosen actions for bad objects in the choice task**

Changes of the proportion of chosen actions toward bad objects in the choice task before and after injection of CPP (NMDA-type receptor antagonist) and NBQX (AMPA-type receptor antagonist) mixture or saline into the GPe (A) and SNr (B). The asterisk indicates a significant difference in proportions of chosen action for Bad objects in comparison among the conditions in the choice tasks (posthoc pairwise t-tests with Bonferroni correction, **p <* 0.05, ***p* < 0.01, ****p* < 0.001, *****p* < 0.0001). Asterisks are attached only to combinations with significant differences.

**Table S1. Summary of statistical test to compare the normalized neuronal activity of STN, GPe, and SNr neurons among conditions during choice task in Figure 2.**

| ***Good-pref STN*** |  |  |  |  |  |  |
| --- | --- | --- | --- | --- | --- | --- |
| parametric bootstrap test (n = 10,000) | *p* |  |  |  |  |  |
| full model vs. null model | < .0001 |  |  |  |  |  |
| post-hoc  (pairwise t-test, Bonferroni correction) | Mean  (SD) | Mean  (SD) | *t* | *p* | 95% CI | effect size |
| good (choice task) vs. good (fixation task) | 1.71  (0.62) | 0.45  (1.00) | 1.28 | < .0001 | [1.01 1.54] | 1.35 |
| good (choice task) vs. bad (choice task) | 1.71  (0.62) | –0.29  (1.41) | 2.00 | < .0001 | [1.79 2.22] | 2.11 |
| good (choice task) vs. bad (fixation task) | 1.71  (0.62) | 0.19  (0.92) | 1.54 | < .0001 | [1.27 1.81] | 1.63 |
| good (fixation task) vs. bad (choice task) | 0.45  (1.00) | –0.29  (1.41) | 0.72 | < .0001 | [0.46 0.99] | 0.76 |
| good (fixation task) vs. bad (fixation task) | 0.45  (1.00) | 0.19  (0.92) | 0.26 | = .07 | [–0.02 0.55] | 0.28 |
| bad (choice task) vs. bad (fixation task) | –0.29  (1.41) | 0.19  (0.92) | ­–0.46 | < .001 | [–0.73 –0.20] | –0.49 |
| ***Bad-pref STN*** |  |  |  |  |  |  |
| parametric bootstrap test (n = 10,000) | *p* |  |  |  |  |  |
| full model vs. null model | < .01 |  |  |  |  |  |
| post-hoc  (pairwise t-test, Bonferroni correction) | Mean  (SD) | Mean  (SD) | *t* | *p* | 95% CI | effect size |
| good (choice task) vs. good (fixation task) | 0.68  (0.99) | 0.71  (1.30) | –0.05 | = .90 | [–0.79 0.70] | –0.05 |
| good (choice task) vs. bad (choice task) | 0.68  (0.99) | 1.50  (0.98) | –0.82 | < .01 | [–1.29 –0.35] | –0.87 |
| good (choice task) vs. bad (fixation task) | 0.68  (0.99) | 0.75  (1.23) | –0.08 | = .83 | [–0.82 0.66] | –0.09 |
| good (fixation task) vs. bad (choice task) | 0.71  (1.30) | 1.50  (0.98) | –0.77 | = .04 | [–1.51 –0.03] | –0.82 |
| good (fixation task) vs. bad (fixation task) | 0.71  (1.30) | 0.75  (1.23) | –0.03 | = .93 | [–0.80 0.73] | –0.04 |
| bad (choice task) vs. bad (fixation task) | 1.50  (0.98) | 0.75  (1.23) | 0.74 | = .05 | [–0.01 1.48] | 0.79 |
| ***Good-pref GPe*** |  |  |  |  |  |  |
| parametric bootstrap test (n = 10,000) | *p* |  |  |  |  |  |
| full model vs. null model | < .0001 |  |  |  |  |  |
| Post-hoc  (pairwise t-test, Bonferroni correction) | Mean  (SD) | Mean  (SD) | *t* | *p* | 95% CI | effect size |
| good (choice task) vs. good (fixation task) | 1.75  (0.76) | 0.80  (1.22) | 0.97 | < .0001 | [0.69 1.24] | 0.93 |
| good (choice task) vs. bad (choice task) | 1.75  (0.76) | –0.10  (1.54) | 1.85 | < .0001 | [1.59 2.11] | 1.77 |
| good (choice task) vs. bad (fixation task) | 1.75  (0.76) | 0.65  (1.19) | 1.12 | < .0001 | [0.84 1.40] | 1.07 |
| good (fixation task) vs. bad (choice task) | 0.80  (1.22) | –0.10  (1.54) | 0.88 | < .0001 | [0.61 1.16] | 0.85 |
| good (fixation task) vs. bad (fixation task) | 0.80  (1.22) | 0.65  (1.19) | 0.16 | = .29 | [–0.13 0.44] | 0.15 |
| bad (choice task) vs. bad (fixation task) | –0.10  (1.54) | 0.65  (1.19) | –0.73 | < .0001 | [–1.01 –0.45] | –0.70 |
| ***Bad-pref GPe*** |  |  |  |  |  |  |
| parametric bootstrap test (n = 10,000) | *p* |  |  |  |  |  |
| full model vs. null model | < .0001 |  |  |  |  |  |
| post-hoc  (pairwise t-test, Bonferroni correction) | Mean  (SD) | Mean  (SD) | *t* | *p* | 95% CI | effect size |
| good (choice task) vs. good (fixation task) | –0.40  (1.24) | –0.16  (1.11) | –0.26 | = .11 | [–0.57 0.06] | –0.26 |
| good (choice task) vs. bad (choice task) | –0.40  (1.24) | –1.57  (0.91) | 1.17 | < .0001 | [0.87 1.48] | 1.20 |
| good (choice task) vs. bad (fixation task) | –0.40  (1.24) | –0.24  (1.11) | –0.18 | = .25 | [–0.50 0.13] | –0.19 |
| good (fixation task) vs. bad (choice task) | –0.16  (1.11) | –1.57  (0.91) | 1.43 | < .0001 | [1.12 1.74] | 1.46 |
| good (fixation task) vs. bad (fixation task) | –0.16  (1.11) | –0.24  (1.11) | 0.07 | = .65 | [–0.25 0.40] | 0.08 |
| bad (choice task) vs. bad (fixation task) | –1.57  (0.91) | –0.24  (1.11) | –1.36 | < .0001 | [–1.67 –1.04] | –1.38 |
| ***SNr*** |  |  |  |  |  |  |
| parametric bootstrap test (n = 10,000) | *p* |  |  |  |  |  |
| full model vs. null model | < .0001 |  |  |  |  |  |
| post-hoc  (pairwise t-test, Bonferroni correction) | Mean  (SD) | Mean  (SD) | *t* | *p* | 95% CI | effect size |
| good (choice task) vs. good (fixation task) | –1.31  (1.02) | 1.22  (1.00) | –2.58 | < .0001 | [–2.84 –2.31] | –2.98 |
| good (choice task) vs. bad (choice task) | –1.31  (1.02) | 1.47  (1.08) | –2.78 | < .0001 | [–3.02 –2.54] | –3.22 |
| good (choice task) vs. bad (fixation task) | –1.31  (1.02) | 1.47  (0.88) | –2.78 | < .0001 | [–3.05 –2.52] | –3.23 |
| good (fixation task) vs. bad (choice task) | 1.22  (1.00) | 1.47  (1.08) | –0.21 | = .13 | [–0.47 0.06] | –0.24 |
| good (fixation task) vs. bad (fixation task) | 1.22  (1.00) | 1.47  (0.88) | –0.21 | = .15 | [–0.49 0.07] | –0.24 |
| bad (choice task) vs. bad (fixation task) | 1.47  (1.08) | 1.47  (0.88) | –0.00 | = .99 | [–0.27 0.27] | –0.00 |

| ***Good-pref STN*** |  |  |  |  |  |  |
| --- | --- | --- | --- | --- | --- | --- |
| parametric bootstrap test (n = 10,000) | *p* |  |  |  |  |  |
| full model vs. null model | < .0001 |  |  |  |  |  |
| post-hoc  (pairwise t-test, Bonferroni correction) | Mean (SD) | Mean (SD) | *t* | *p* | 95% CI | effect size |
| **scene1** |  |  |  |  |  |  |
| (good, contra) vs (bad, contra) | 1.71 (0.62) | –0.29 (1.41) | 2.00 | < .0001 | [1.82 2.18] | 2.49 |
| (good, contra) vs (good, ipsi) | 1.71 (0.62) | 0.83 (1.07) | 0.88 | < .0001 | [0.70 1.07] | 1.10 |
| (good, contra) vs (bad, ipsi) | 1.71 (0.62) | –0.45 (1.29) | 2.16 | < .0001 | [1.98 2.34] | 2.69 |
| (bad, contra) vs (good, ipsi) | –0.29 (1.41) | 0.83 (1.07) | –1.12 | < .0001 | [–1.30 –0.94] | –1.39 |
| (bad, contra) vs (bad, ipsi) | –0.29 (1.41) | –0.45 (1.29) | 0.16 | = .08 | [–0.02 0.34] | 0.20 |
| (good, ipsi) vs (bad, ipsi) | 0.83 (1.07) | –0.45 (1.29) | 1.28 | < .0001 | [1.10 1.46] | 1.59 |
| **scene2** |  |  |  |  |  |  |
| (good, contra) vs (bad, contra) | 1.73 (0.61) | –0.33 (1.23) | 2.06 | < .0001 | [1.88 2.24] | 2.57 |
| (good, contra) vs (good, ipsi) | 1.73 (0.61) | 0.87 (0.97) | 0.86 | < .0001 | [0.68 1.04] | 1.07 |
| (good, contra) vs (bad, ipsi) | 1.73 (0.61) | –0.52 (1.20) | 2.25 | < .0001 | [2.06 2.43] | 2.80 |
| (bad, contra) vs (good, ipsi) | –0.33 (1.23) | 0.87 (0.97) | –1.20 | < .0001 | [–1.38 –1.02] | –1.50 |
| (bad, contra) vs (bad, ipsi) | –0.33 (1.23) | –0.52 (1.20) | 0.18 | = .05 | [0.00 0.37] | 0.23 |
| (good, ipsi) vs (bad, ipsi) | 0.87 (0.97) | –0.52 (1.20) | 1.39 | < .0001 | [1.20 1.57] | 1.73 |
| **scene3** |  |  |  |  |  |  |
| (good, contra) vs (bad, contra) | 1.75 (0.60) | –0.10 (1.31) | 1.85 | < .0001 | [1.67 2.03] | 2.30 |
| (good, contra) vs (good, ipsi) | 1.75 (0.60) | 0.91 (1.01) | 0.84 | < .0001 | [0.66 1.02] | 1.04 |
| (good, contra) vs (bad, ipsi) | 1.75 (0.60) | –0.42 (1.28) | 2.17 | < .0001 | [1.99 2.35] | 2.70 |
| (bad, contra) vs (good, ipsi) | –0.10 (1.31) | 0.91 (1.01) | –1.01 | < .0001 | [–1.19 –0.83] | –1.26 |
| (bad, contra) vs (bad, ipsi) | –0.10 (1.31) | –0.42 (1.28) | 0.32 | < .001 | [0.14 0.50] | 0.40 |
| (good, ipsi) vs (bad, ipsi) | 0.91 (1.01) | –0.42 (1.28) | 1.33 | < .0001 | [1.15 1.51] | 1.66 |
| **scene4** |  |  |  |  |  |  |
| (good, contra) vs (bad, contra) | 1.73 (0.59) | –0.10 (1.33) | 1.84 | < .0001 | [1.66 2.02] | 2.29 |
| (good, contra) vs (good, ipsi) | 1.73 (0.59) | 0.83 (1.03) | 0.90 | < .0001 | [0.72 1.08] | 1.13 |
| (good, contra) vs (bad, ipsi) | 1.73 (0.59) | –0.39 (1.27) | 1.22 | < .0001 | [1.94 2.30] | 2.64 |
| (bad, contra) vs (good, ipsi) | –0.10 (1.33) | 0.83 (1.03) | –0.93 | < .0001 | [–1.12 –0.75] | –1.16 |
| (bad, contra) vs (bad, ipsi) | –0.10 (1.33) | –0.39 (1.27) | 1.22 | < .01 | [0.10 0.47] | 0.35 |
| (good, ipsi) vs (bad, ipsi) | 0.83 (1.03) | –0.39 (1.27) | 1.22 | < .0001 | [1.04 1.40] | 1.52 |

**Table S2. Summary of statistical test to compare the normalized neuronal activity of Good-pref STN neurons at target onset among conditions during choice task in Figure S4.**

| ***Bad-pref STN*** |  |  |  |  |  |  |
| --- | --- | --- | --- | --- | --- | --- |
| parametric bootstrap test (n = 10,000) | *p* |  |  |  |  |  |
| full model vs. null model | < .0001 |  |  |  |  |  |
| post-hoc  (pairwise t-test, Bonferroni correction) | Mean (SD) | Mean (SD) | *t* | *p* | 95% CI | effect size |
| **scene1** |  |  |  |  |  |  |
| (good, contra) vs (bad, contra) | 0.68 (0.99) | 1.50 (0.98) | –0.82 | < .0001 | [–1.20 –0.44] | –1.06 |
| (good, contra) vs (good, ipsi) | 0.68 (0.99) | 0.11 (1.15) | 0.58 | < .01 | [0.20 0.96] | 0.75 |
| (good, contra) vs (bad, ipsi) | 0.68 (0.99) | 1.46 (0.93) | –0.77 | = .0001 | [–1.15 –0.40] | –1.00 |
| (bad, contra) vs (good, ipsi) | 1.50 (0.98) | 0.11 (1.15) | 1.39 | < .0001 | [1.01 1.77] | 1.80 |
| (bad, contra) vs (bad, ipsi) | 1.50 (0.98) | 1.46 (0.93) | 0.04 | = .83 | [–0.34 0.42] | 0.05 |
| (good, ipsi) vs (bad, ipsi) | 0.11 (1.15) | 1.46 (0.93) | –1.35 | < .0001 | [–1.73 –0.97] | –1.75 |
| **scene2** |  |  |  |  |  |  |
| (good, contra) vs (bad, contra) | 0.44 (1.21) | 1.63 (0.93) | –1.19 | < .0001 | [–1.56 –0.81] | –1.53 |
| (good, contra) vs (good, ipsi) | 0.44 (1.21) | 0.25 (1.27) | 0.19 | = .32 | [–0.19 0.57] | 0.25 |
| (good, contra) vs (bad, ipsi) | 0.44 (1.21) | 1.40 (1.03) | –0.96 | < .0001 | [–1.34 –0.58] | –1.24 |
| (bad, contra) vs (good, ipsi) | 1.63 (0.93) | 0.25 (1.27) | 1.38 | < .0001 | [1.00 1.76] | 1.79 |
| (bad, contra) vs (bad, ipsi) | 1.63 (0.93) | 1.40 (1.03) | 0.23 | = .24 | [–0.15 0.61] | 0.30 |
| (good, ipsi) vs (bad, ipsi) | 0.25 (1.27) | 1.40 (1.03) | –1.15 | < .0001 | [–1.53 –0.77] | –1.49 |
| **scene3** |  |  |  |  |  |  |
| (good, contra) vs (bad, contra) | 0.53 (1.19) | 1.62 (0.95) | –1.09 | < .0001 | [–1.47 –0.71] | –1.41 |
| (good, contra) vs (good, ipsi) | 0.53 (1.19) | 0.17 (1.15) | 0.36 | = .0598 | [-0.02 0.74] | 0.47 |
| (good, contra) vs (bad, ipsi) | 0.53 (1.19) | 1.39 (0.95) | –0.86 | < .0001 | [0.48 1.23] | –1.11 |
| (bad, contra) vs (good, ipsi) | 1.62 (0.95) | 0.17 (1.15) | 1.46 | < .0001 | [1.08 1.84] | 1.88 |
| (bad, contra) vs (bad, ipsi) | 1.62 (0.95) | 1.39 (0.95) | 0.23 | = .228 | –0.15 0.61] | 0.30 |
| (good, ipsi) vs (bad, ipsi) | 0.17 (1.15) | 1.39 (0.95) | –1.22 | < .0001 | [–1.60 –0.84] | –1.58 |
| **scene4** |  |  |  |  |  |  |
| (good, contra) vs (bad, contra) | 0.65 (0.98) | 1.50 (0.96) | –0.85 | < .0001 | [–1.23 –0.47] | –1.10 |
| (good, contra) vs (good, ipsi) | 0.65 (0.98) | 0.23 (1.19) | 0.42 | < .05 | [0.04 0.80] | 0.55 |
| (good, contra) vs (bad, ipsi) | 0.65 (0.98) | 1.29 (1.07) | –0.64 | = .001 | [–1.02 –0.26] | –0.83 |
| (bad, contra) vs (good, ipsi) | 1.50 (0.96) | 0.23 (1.19) | 1.27 | < .0001 | [0.89 1.65] | 1.65 |
| (bad, contra) vs (bad, ipsi) | 1.50 (0.96) | 1.29 (1.07) | 0.21 | = .277 | [–0.17 0.59] | 0.27 |
| (good, ipsi) vs (bad, ipsi) | 0.23 (1.19) | 1.29 (1.07) | –1.06 | < .0001 | [–1.44 –0.68] | –1.38 |

**Table S3. Summary of statistical test to compare the normalized neuronal activity of Bad-pref STN neurons at target onset among conditions during choice task in Figure S4.**

| ***Good-pref GPe*** |  |  |  |  |  |  |
| --- | --- | --- | --- | --- | --- | --- |
| parametric bootstrap test (n = 10,000) | *p* |  |  |  |  |  |
| full model vs. null model | < .0001 |  |  |  |  |  |
| post-hoc  (pairwise t-test, Bonferroni correction) | Mean (SD) | Mean (SD) | *t* | *p* | 95% CI | effect size |
| **scene1** |  |  |  |  |  |  |
| (good, contra) vs (bad, contra) | 1.75 (0.76) | –0.10 (1.54) | 1.85 | < .0001 | [1.61 2.09] | 1.95 |
| (good, contra) vs (good, ipsi) | 1.75 (0.76) | 0.80 (0.98) | 0.94 | < .0001 | [0.70 1.19] | 0.98 |
| (good, contra) vs (bad, ipsi) | 1.75 (0.76) | –0.40 (1.51) | 2.14 | < .0001 | [1.90 2.38] | 2.22 |
| (bad, contra) vs (good, ipsi) | –0.10 (1.54) | 0.80 (0.98) | –0.90 | < .0001 | [–1.14 –0.66] | –0.94 |
| (bad, contra) vs (bad, ipsi) | –0.10 (1.54) | –0.40 (1.51) | 0.29 | < .05 | [ 0.05 0.53] | 0.30 |
| (good, ipsi) vs (bad, ipsi) | 0.80 (0.98) | –0.40 (1.51) | 1.19 | < .0001 | [0.95 1.43] | 1.24 |
| **scene2** |  |  |  |  |  |  |
| (good, contra) vs (bad, contra) | 1.75 (0.86) | –0.13 (1.47) | 1.88 | < .0001 | [1.64 2.12] | 1.95 |
| (good, contra) vs (good, ipsi) | 1.75 (0.86) | 0.97 (1.09) | 0.79 | < .0001 | [0.54 1.03] | 0.81 |
| (good, contra) vs (bad, ipsi) | 1.75 (0.86) | –0.32 (1.34) | 2.09 | < .0001 | [1.85 2.33] | 2.16 |
| (bad, contra) vs (good, ipsi) | –0.13 (1.47) | 0.97 (1.09) | –1.10 | < .0001 | [–1.34 –0.86] | –1.14 |
| (bad, contra) vs (bad, ipsi) | –0.13 (1.47) | –0.32 (1.34) | 0.21 | = .10 | [–0.04 0.45] | 0.21 |
| (good, ipsi) vs (bad, ipsi) | 0.97 (1.09) | –0.32 (1.34) | 1.30 | < .0001 | [1.06 1.54] | 1.35 |
| **scene3** |  |  |  |  |  |  |
| (good, contra) vs (bad, contra) | 1.77 (0.81) | 0.19 (1.08) | 1.58 | < .0001 | [1.34 1.82] | 1.64 |
| (good, contra) vs (good, ipsi) | 1.77 (0.81) | 0.94 (1.08) | 0.83 | < .0001 | [0.59 1.07] | 0.86 |
| (good, contra) vs (bad, ipsi) | 1.77 (0.81) | –0.20 (1.37) | 1.97 | < .0001 | [1.73 2.21] | 2.05 |
| (bad, contra) vs (good, ipsi) | 0.19 (1.08) | 0.94 (1.08) | –0.75 | < .0001 | [–0.99 –0.51] | –0.78 |
| (bad, contra) vs (bad, ipsi) | 0.19 (1.08) | –0.20 (1.37) | 0.39 | < .01 | [0.15 0.63] | 0.40 |
| (good, ipsi) vs (bad, ipsi) | 0.94 (1.08) | –0.20 (1.37) | 1.14 | < .0001 | [0.90 1.38] | 1.18 |
| **scene4** |  |  |  |  |  |  |
| (good, contra) vs (bad, contra) | 1.66 (0.87) | –0.08 (1.75) | 1.74 | < .0001 | [1.50 1.98] | 1.81 |
| (good, contra) vs (good, ipsi) | 1.66 (0.87) | 0.82 (1.11) | 0.84 | < .0001 | [0.60 1.08] | 0.87 |
| (good, contra) vs (bad, ipsi) | 1.66 (0.87) | –0.47 (1.48) | 2.13 | < .0001 | [1.89 2.37] | 2.21 |
| (bad, contra) vs (good, ipsi) | –0.08 (1.75) | 0.82 (1.11) | –0.90 | < .0001 | [–1.14 –0.66] | –0.94 |
| (bad, contra) vs (bad, ipsi) | –0.08 (1.75) | –0.47 (1.48) | 0.38 | < .01 | [0.14 0.63] | 0.40 |
| (good, ipsi) vs (bad, ipsi) | 0.82 (1.11) | –0.47 (1.48) | 1.29 | < .0001 | [1.05 1.53] | 1.34 |

**Table S4. Summary of statistical test to compare the normalized neuronal activity of Good-pref GPe neurons at target onset among conditions during choice task in Figure S3.**

| ***Bad-pref GPe*** |  |  |  |  |  |  |
| --- | --- | --- | --- | --- | --- | --- |
| parametric bootstrap test (n = 10,000) | *p* |  |  |  |  |  |
| full model vs. null model | < .0001 |  |  |  |  |  |
| post-hoc  (pairwise t-test, Bonferroni correction) | Mean (SD) | Mean (SD) | *t* | *p* | 95% CI | effect size |
| **scene1** |  |  |  |  |  |  |
| (good, contra) vs (bad, contra) | –0.42 (1.24) | –1.58 (0.91) | 1.17 | < .0001 | [0.95 1.39] | 1.65 |
| (good, contra) vs (good, ipsi) | –0.42 (1.24) | –0.85 (1.12) | 0.45 | < .001 | [0.23 0.67] | 0.63 |
| (good, contra) vs (bad, ipsi) | –0.42 (1.24) | –1.60 (0.86) | 1.19 | < .0001 | [0.98 1.41] | 1.68 |
| (bad, contra) vs (good, ipsi) | –1.58 (0.91) | –0.85 (1.12) | –0.72 | < .0001 | [–0.94 –0.50] | –1.06 |
| (bad, contra) vs (bad, ipsi) | –1.58 (0.91) | –1.60 (0.86) | 0.22 | = .84 | [–0.20 0.24] | 0.03 |
| (good, ipsi) vs (bad, ipsi) | –0.85 (1.12) | –1.60 (0.86) | 0.75 | < .0001 | [0.53 0.96] | 1.05 |
| **scene2** |  |  |  |  |  |  |
| (good, contra) vs (bad, contra) | –0.63 (1.21) | –1.60 (0.82) | 0.97 | < .0001 | [0.75 1.19] | 1.36 |
| (good, contra) vs (good, ipsi) | –0.63 (1.21) | –0.99 (1.14) | 0.36 | < .01 | [0.14 0.58] | 0.50 |
| (good, contra) vs (bad, ipsi) | –0.63 (1.21) | –1.73 (0.81) | 1.10 | < .0001 | [0.88 1.32] | 1.54 |
| (bad, contra) vs (good, ipsi) | –1.60 (0.82) | –0.99 (1.14) | –0.61 | < .0001 | [–0.83 –0.39] | –0.86 |
| (bad, contra) vs (bad, ipsi) | –1.60 (0.82) | –1.73 (0.81) | 0.13 | = .26 | [–0.09 0.35] | 0.18 |
| (good, ipsi) vs (bad, ipsi) | –0.99 (1.14) | –1.73 (0.81) | 0.74 | < .0001 | [0.52 0.96] | 1.04 |
| **scene3** |  |  |  |  |  |  |
| (good, contra) vs (bad, contra) | –0.58 (1.15) | –1.66 (0.83) | 1.09 | < .0001 | [0.87 1.31] | 1.53 |
| (good, contra) vs (good, ipsi) | –0.58 (1.15) | –0.98 (1.17) | 0.41 | < .001 | [0.19 0.63] | 0.57 |
| (good, contra) vs (bad, ipsi) | –0.58 (1.15) | –1.73 (0.84) | 1.15 | < .0001 | [0.94 1.37] | 1.62 |
| (bad, contra) vs (good, ipsi) | –1.66 (0.83) | –0.98 (1.17) | –0.68 | < .0001 | [–0.90 –0.46] | –0.96 |
| (bad, contra) vs (bad, ipsi) | –1.66 (0.83) | –1.73 (0.84) | 0.69 | = .54 | [–0.15 0.29] | 0.10 |
| (good, ipsi) vs (bad, ipsi) | –0.98 (1.17) | –1.73 (0.84) | 0.75 | < .0001 | [0.53 0.97] | 1.05 |
| **scene4** |  |  |  |  |  |  |
| (good, contra) vs (bad, contra) | –0.50 (1.28) | –1.54 (0.88) | 1.04 | < .0001 | [0.82 1.26] | 1.46 |
| (good, contra) vs (good, ipsi) | –0.50 (1.28) | –0.87 (1.17) | 0.37 | < .001 | [0.15 0.59] | 0.52 |
| (good, contra) vs (bad, ipsi) | –0.50 (1.28) | –1.51 (0.84) | 1.01 | < .0001 | [0.79 1.23] | 1.42 |
| (bad, contra) vs (good, ipsi) | –1.54 (0.88) | –0.87 (1.17) | –0.66 | < .0001 | [–0.88 –0.44] | –0.93 |
| (bad, contra) vs (bad, ipsi) | –1.54 (0.88) | –1.51 (0.84) | –0.03 | = .82 | [–0.25 0.19] | –0.04 |
| (good, ipsi) vs (bad, ipsi) | –0.87 (1.17) | –1.51 (0.84) | 0.64 | < .0001 | [0.42 0.86] | 0.90 |

**Table S5. Summary of statistical test to compare the normalized neuronal activity of Bad-pref GPe neurons at target onset among conditions during choice task in Figure S3.**

| ***SNr*** |  |  |  |  |  |  |
| --- | --- | --- | --- | --- | --- | --- |
| parametric bootstrap test (n = 10,000) | *p* |  |  |  |  |  |
| full model vs. null model | < .0001 |  |  |  |  |  |
| post-hoc  (pairwise t-test, Bonferroni correction) | Mean (SD) | Mean (SD) | *t* | *p* | 95% CI | effect size |
| **scene1** |  |  |  |  |  |  |
| (good, contra) vs (bad, contra) | –1.31 (1.02) | 1.47 (1.08) | –2.79 | < .0001 | [–3.02 –2.56] | –3.37 |
| (good, contra) vs (good, ipsi) | –1.31 (1.02) | –0.98 (1.08) | –0.34 | < .01 | [–0.57 –0.11] | –0.41 |
| (good, contra) vs (bad, ipsi) | –1.31 (1.02) | 1.07 (1.21) | –2.38 | < .0001 | [–2.61 –2.15] | –2.88 |
| (bad, contra) vs (good, ipsi) | 1.47 (1.08) | –0.98 (1.08) | 2.45 | < .0001 | [2.22 2.68] | 2.96 |
| (bad, contra) vs (bad, ipsi) | 1.47 (1.08) | 1.07 (1.21) | 0.40 | < .001 | [0.17 0.63] | 0.49 |
| (good, ipsi) vs (bad, ipsi) | –0.98 (1.08) | 1.07 (1.21) | –2.05 | < .0001 | [–2.27 –1.82] | –2.47 |
| **scene2** |  |  |  |  |  |  |
| (good, contra) vs (bad, contra) | –1.30 (1.09) | 1.71 (0.88) | –3.02 | < .0001 | [–3.24 –2.78] | –3.64 |
| (good, contra) vs (good, ipsi) | –1.30 (1.09) | –0.88 (1.05) | –0.43 | < .001 | [–0.66 –0.20] | –0.52 |
| (good, contra) vs (bad, ipsi) | –1.30 (1.09) | 1.28 (1.04) | –2.58 | < .0001 | [–2.81 –2.36] | –3.13 |
| (bad, contra) vs (good, ipsi) | 1.71 (0.88) | –0.88 (1.05) | 2.59 | < .0001 | [2.36 2.82] | 3.13 |
| (bad, contra) vs (bad, ipsi) | 1.71 (0.88) | 1.28 (1.04) | 0.43 | < .001 | [0.20 0.66] | 0.52 |
| (good, ipsi) vs (bad, ipsi) | –0.88 (1.05) | 1.28 (1.04) | –2.16 | < .0001 | [–2.39 –1.93] | –2.61 |
| **scene3** |  |  |  |  |  |  |
| (good, contra) vs (bad, contra) | –1.56 (0.88) | 1.55 (1.05) | –3.11 | < .0001 | [–3.34 –2.88] | –3.76 |
| (good, contra) vs (good, ipsi) | –1.56 (0.88) | –1.11 (1.01) | –0.44 | < .001 | [–0.67 –0.21] | –0.53 |
| (good, contra) vs (bad, ipsi) | –1.56 (0.88) | 1.32 (0.94) | –2.88 | < .0001 | [–3.10 –2.65] | –3.48 |
| (bad, contra) vs (good, ipsi) | 1.55 (1.05) | –1.11 (1.01) | 2.67 | < .0001 | [2.44 2.90] | 3.23 |
| (bad, contra) vs (bad, ipsi) | 1.55 (1.05) | 1.32 (0.94) | 0.24 | < .05 | [0.01 0.47] | 0.29 |
| (good, ipsi) vs (bad, ipsi) | –1.11 (1.01) | 1.32 (0.94) | –2.43 | < .0001 | [–2.66 –2.20] | –2.94 |
| **scene4** |  |  |  |  |  |  |
| (good, contra) vs (bad, contra) | –1.34 (1.14) | 1.61 (0.99) | –2.95 | < .0001 | [–3.18 –2.72] | –3.57 |
| (good, contra) vs (good, ipsi) | –1.34 (1.14) | –0.90 (1.11) | –0.44 | < .001 | [-0.67 –0.21] | –0.53 |
| (good, contra) vs (bad, ipsi) | –1.34 (1.14) | 1.30 (1.08) | –2.64 | < .0001 | [–2.86 –2.41] | –3.19 |
| (bad, contra) vs (good, ipsi) | 1.61 (0.99) | –0.90 (1.11) | 2.51 | < .0001 | [2.28 2.74] | 3.04 |
| (bad, contra) vs (bad, ipsi) | 1.61 (0.99) | 1.30 (1.08) | 0.31 | < .01 | [0.08 0.54] | 0.38 |
| (good, ipsi) vs (bad, ipsi) | –0.90 (1.11) | 1.30 (1.08) | –2.20 | < .0001 | [–2.43 –1.97] | –2.66 |

**Table S6. Summary of statistical test to compare the normalized neuronal activity of SNr neurons at target onset among conditions during choice task in Figure S3.**

| ***Good-pref STN*** |  |  |  |  |  |  |
| --- | --- | --- | --- | --- | --- | --- |
| parametric bootstrap test (n = 10,000) | *p* |  |  |  |  |  |
| full model vs. null model | < .0001 |  |  |  |  |  |
| post-hoc  (pairwise t-test, Bonferroni correction) | Mean (SD) | Mean (SD) | *t* | *p* | 95% CI | effect size |
| **scene1** |  |  |  |  |  |  |
| (good, contra) vs (bad, contra) | 1.53 (0.56) | –0.37 (1.43) | 1.90 | < .0001 | [1.73 2.08] | 2.44 |
| (good, contra) vs (good, ipsi) | 1.53 (0.56) | 0.82 (0.87) | 0.72 | < .0001 | [0.54 0.90] | 0.92 |
| (good, contra) vs (bad, ipsi) | 1.53 (0.56) | –0.54 (1.24) | 2.08 | < .0001 | [1.90 2.25] | 2.66 |
| (bad, contra) vs (good, ipsi) | –0.37 (1.43) | 0.82 (0.87) | –1.19 | < .0001 | [–1.36 –1.00] | –1.52 |
| (bad, contra) vs (bad, ipsi) | –0.37 (1.43) | –0.54 (1.24) | 0.17 | = .053 | [–0.00 0.35] | 0.22 |
| (good, ipsi) vs (bad, ipsi) | 0.82 (0.87) | –0.54 (1.24) | 1.36 | < .0001 | [1.18 1.54] | 1.74 |
| **scene2** |  |  |  |  |  |  |
| (good, contra) vs (bad, contra) | 1.59 (0.58) | –0.47 (1.31) | 2.06 | < .0001 | [1.88 2.24] | 2.64 |
| (good, contra) vs (good, ipsi) | 1.59 (0.58) | 0.88 (0.79) | 0.71 | < .0001 | [0.53 0.88] | 0.91 |
| (good, contra) vs (bad, ipsi) | 1.59 (0.58) | –0.61 (1.25) | 2.20 | < .0001 | [2.02 2.37] | 2.81 |
| (bad, contra) vs (good, ipsi) | –0.47 (1.31) | 0.88 (0.79) | –1.35 | < .0001 | [–1.53 –1.18] | –1.73 |
| (bad, contra) vs (bad, ipsi) | –0.47 (1.31) | –0.61 (1.25) | 0.14 | = .134 | [–0.04 0.31] | 0.17 |
| (good, ipsi) vs (bad, ipsi) | 0.88 (0.79) | –0.61 (1.25) | 1.49 | < .0001 | [1.31 1.67] | 1.91 |
| **scene3** |  |  |  |  |  |  |
| (good, contra) vs (bad, contra) | 1.56 (0.62) | –0.09 (1.32) | 1.65 | < .0001 | [1.47 1.83] | 2.11 |
| (good, contra) vs (good, ipsi) | 1.56 (0.62) | 0.91 (0.86) | 0.65 | < .0001 | [0.47 0.83] | 0.83 |
| (good, contra) vs (bad, ipsi) | 1.56 (0.62) | –0.38 (1.30) | 1.94 | < .0001 | [1.76 2.12] | 2.49 |
| (bad, contra) vs (good, ipsi) | –0.09 (1.32) | 0.91 (0.86) | –1.00 | < .0001 | [–1.18 –0.83] | –1.28 |
| (bad, contra) vs (bad, ipsi) | –0.09 (1.32) | –0.38 (1.30) | 0.29 | < .01 | [0.11 0.47] | 0.37 |
| (good, ipsi) vs (bad, ipsi) | 0.91 (0.86) | –0.38 (1.30) | 1.29 | < .0001 | [1.12 1.47] | 1.66 |
| **scene4** |  |  |  |  |  |  |
| (good, contra) vs (bad, contra) | 1.58 (0.54) | –0.15 (1.37) | 1.73 | < .0001 | [1.55 1.91] | 2.22 |
| (good, contra) vs (good, ipsi) | 1.58 (0.54) | 0.83 (0.92) | 0.75 | < .0001 | [0.58 0.93] | 0.96 |
| (good, contra) vs (bad, ipsi) | 1.58 (0.54) | –0.45 (1.31) | 2.03 | < .0001 | [1.85 2.20] | 2.59 |
| (bad, contra) vs (good, ipsi) | –0.15 (1.37) | 0.83 (0.92) | –0.98 | < .0001 | [–1.15 –0.80] | –1.25 |
| (bad, contra) vs (bad, ipsi) | –0.15 (1.37) | –0.45 (1.31) | 0.30 | < .01 | [0.12 0.47] | 0.38 |
| (good, ipsi) vs (bad, ipsi) | 0.83 (0.92) | –0.45 (1.31) | 1.27 | < .0001 | [1.10 1.45] | 1.63 |

**Table S7. Summary of statistical test to compare the normalized neuronal activity of Good-pref STN neurons at saccade onset among conditions during choice task in Figure S4.**

| ***Bad-pref STN*** |  |  |  |  |  |  |
| --- | --- | --- | --- | --- | --- | --- |
| parametric bootstrap test (n = 10,000) | *p* |  |  |  |  |  |
| full model vs. null model | < .0001 |  |  |  |  |  |
| post-hoc  (pairwise t-test, Bonferroni correction) | Mean (SD) | Mean (SD) | *t* | *p* | 95% CI | effect size |
| **scene1** |  |  |  |  |  |  |
| (good, contra) vs (bad, contra) | 0.84 (0.94) | 1.46 (0.90) | –0.62 | < .001 | [–0.96 –0.28] | –0.90 |
| (good, contra) vs (good, ipsi) | 0.84 (0.94) | 0.36 (1.12) | 0.48 | < .01 | [0.14 0.81] | 0.70 |
| (good, contra) vs (bad, ipsi) | 0.84 (0.94) | 1.38 (0.96) | –0.55 | < .01 | [–0.88 –0.21] | –0.80 |
| (bad, contra) vs (good, ipsi) | 1.46 (0.90) | 0.36 (1.12) | 1.10 | < .0001 | [0.76 1.43] | 1.60 |
| (bad, contra) vs (bad, ipsi) | 1.46 (0.90) | 1.38 (0.96) | 0.07 | = .66 | [–0.26 0.41] | 0.12 |
| (good, ipsi) vs (bad, ipsi) | 0.36 (1.12) | 1.38 (0.96) | –1.02 | < .0001 | [–1.36 –0.69] | –1.49 |
| **scene2** |  |  |  |  |  |  |
| (good, contra) vs (bad, contra) | 0.65 (1.13) | 1.64 (0.82) | –1.00 | < .0001 | [–1.33 –0.66] | –1.45 |
| (good, contra) vs (good, ipsi) | 0.65 (1.13) | 0.49 (1.12) | 0.16 | = .35 | [–0.18 0.50] | 0.23 |
| (good, contra) vs (bad, ipsi) | 0.65 (1.13) | 1.45 (1.06) | –0.80 | < .0001 | [–1.14 –0.47] | –1.17 |
| (bad, contra) vs (good, ipsi) | 1.64 (0.82) | 0.49 (1.12) | 1.15 | < .0001 | [ 0.82 1.49] | 1.68 |
| (bad, contra) vs (bad, ipsi) | 1.64 (0.82) | 1.45 (1.06) | 0.19 | = .26 | [–0.14 0.53] | 0.28 |
| (good, ipsi) vs (bad, ipsi) | 0.49 (1.12) | 1.45 (1.06) | –0.96 | < .0001 | [–1.30 –0.62] | –1.40 |
| **scene3** |  |  |  |  |  |  |
| (good, contra) vs (bad, contra) | 0.70 (1.13) | 1.58 (0.85) | –0.88 | < .0001 | [–1.21 –0.54] | –1.28 |
| (good, contra) vs (good, ipsi) | 0.70 (1.13) | 0.37 (0.99) | 0.37 | = .03 | [0.03 0.70] | 0.53 |
| (good, contra) vs (bad, ipsi) | 0.70 (1.13) | 1.29 (0.92) | –0.59 | < .001 | [–0.93 –0.26] | –0.86 |
| (bad, contra) vs (good, ipsi) | 1.58 (0.85) | 0.37 (0.99) | 1.24 | < .0001 | [0.91 1.58] | 1.81 |
| (bad, contra) vs (bad, ipsi) | 1.58 (0.85) | 1.29 (0.92) | 0.28 | = .10 | [–0.05 0.62] | 0.42 |
| (good, ipsi) vs (bad, ipsi) | 0.37 (0.99) | 1.29 (0.92) | –0.96 | < .0001 | [–1.29 –0.62] | –1.40 |
| **scene4** |  |  |  |  |  |  |
| (good, contra) vs (bad, contra) | 0.74 (0.80) | 1.49 (0.91) | –0.75 | < .0001 | [–1.08 –0.41] | –1.09 |
| (good, contra) vs (good, ipsi) | 0.74 (0.80) | 0.46 (1.09) | 0.28 | = .11 | [–0.06 0.62] | 0.41 |
| (good, contra) vs (bad, ipsi) | 0.74 (0.80) | 1.25 (1.08) | –0.51 | < .01 | [–0.85 –0.17] | –0.74 |
| (bad, contra) vs (good, ipsi) | 1.49 (0.91) | 0.46 (1.09) | 1.02 | < .0001 | [0.69 1.36] | 1.50 |
| (bad, contra) vs (bad, ipsi) | 1.49 (0.91) | 1.25 (1.08) | 0.23 | = .17 | [–0.10 0.58] | 0.35 |
| (good, ipsi) vs (bad, ipsi) | 0.46 (1.09) | 1.25 (1.08) | –0.79 | < .0001 | [–1.12 –0.45] | –1.15 |

**Table S8. Summary of statistical test to compare the normalized neuronal activity of Bad-pref STN neurons at saccade onset among conditions during choice task in Figure S4.**

| ***Good-pref GPe*** |  |  |  |  |  |  |
| --- | --- | --- | --- | --- | --- | --- |
| parametric bootstrap test (n = 10,000) | *p* |  |  |  |  |  |
| full model vs. null model | < .0001 |  |  |  |  |  |
| post-hoc  (pairwise t-test, Bonferroni correction) | Mean (SD) | Mean (SD) | *t* | *p* | 95% CI | effect size |
| **scene1** |  |  |  |  |  |  |
| (good, contra) vs (bad, contra) | 1.37 (0.78) | –0.20 (1.52) | 1.57 | < .0001 | [1.32 1.82] | 1.56 |
| (good, contra) vs (good, ipsi) | 1.37 (0.78) | 0.52 (0.97) | 0.85 | < .0001 | [0.60 1.10] | 0.85 |
| (good, contra) vs (bad, ipsi) | 1.37 (0.78) | –0.46 (1.54) | 1.83 | < .0001 | [1.58 2.08] | 1.82 |
| (bad, contra) vs (good, ipsi) | –0.20 (1.52) | 0.52 (0.97) | –0.72 | < .0001 | [–0.97 –0.47] | –0.71 |
| (bad, contra) vs (bad, ipsi) | –0.20 (1.52) | –0.46 (1.54) | 2.7 | < .05 | [0.01 0.52] | 0.26 |
| (good, ipsi) vs (bad, ipsi) | 0.52 (0.97) | –0.46 (1.54) | 0.98 | < .0001 | [0.73 1.23] | 0.98 |
| **scene2** |  |  |  |  |  |  |
| (good, contra) vs (bad, contra) | 1.45 (0.77) | –0.26 (1.49) | 1.71 | < .0001 | [1.45 1.96] | 1.69 |
| (good, contra) vs (good, ipsi) | 1.45 (0.77) | 0.80 (1.07) | 0.65 | < .0001 | [0.40 0.90] | 0.64 |
| (good, contra) vs (bad, ipsi) | 1.45 (0.77) | –0.36 (1.36) | 1.82 | < .0001 | [1.56 2.07] | 1.80 |
| (bad, contra) vs (good, ipsi) | –0.26 (1.49) | 0.80 (1.07) | –1.06 | < .0001 | [–1.31 –0.81] | –1.05 |
| (bad, contra) vs (bad, ipsi) | –0.26 (1.49) | –0.36 (1.36) | 0.11 | = .385 | [–0.14 0.36] | 0.11 |
| (good, ipsi) vs (bad, ipsi) | 0.80 (1.07) | –0.36 (1.36) | 1.17 | < .0001 | [0.92 1.42] | 1.16 |
| **scene3** |  |  |  |  |  |  |
| (good, contra) vs (bad, contra) | 1.49 (0.74) | 0.12 (1.53) | 1.37 | < .0001 | [1.12 1.62] | 1.36 |
| (good, contra) vs (good, ipsi) | 1.49 (0.74) | 0.77 (1.06) | 0.72 | < .0001 | [0.47 0.97] | 0.72 |
| (good, contra) vs (bad, ipsi) | 1.49 (0.74) | –0.21 (1.38) | 1.70 | < .0001 | [1.45 1.95] | 1.69 |
| (bad, contra) vs (good, ipsi) | 0.12 (1.53) | 0.77 (1.06) | –0.65 | < .0001 | [–0.90 –0.40] | –0.64 |
| (bad, contra) vs (bad, ipsi) | 0.12 (1.53) | –0.21 (1.38) | 0.33 | < .01 | [0.08 0.59] | 0.33 |
| (good, ipsi) vs (bad, ipsi) | 0.77 (1.06) | –0.21 (1.38) | 0.98 | < .0001 | [0.73 1.23] | 0.97 |
| **scene4** |  |  |  |  |  |  |
| (good, contra) vs (bad, contra) | 1.29 (0.83) | –0.14 (1.90) | 1.43 | < .0001 | [1.18 1.68] | 1.42 |
| (good, contra) vs (good, ipsi) | 1.29 (0.83) | 0.57 (1.11) | 0.73 | < .0001 | [0.47 0.98] | 0.72 |
| (good, contra) vs (bad, ipsi) | 1.29 (0.83) | –0.54 (1.53) | 1.83 | < .0001 | [1.58 2.08] | 1.81 |
| (bad, contra) vs (good, ipsi) | –0.14 (1.90) | 0.57 (1.11) | –0.71 | < .0001 | [–0.96 –0.46] | –0.70 |
| (bad, contra) vs (bad, ipsi) | –0.14 (1.90) | –0.54 (1.53) | 0.40 | < .01 | [0.14 0.65] | 0.39 |
| (good, ipsi) vs (bad, ipsi) | 0.57 (1.11) | –0.54 (1.53) | 1.10 | < .0001 | [0.85 1.35] | 1.09 |

**Table S9. Summary of statistical test to compare the normalized neuronal activity of Good-pref GPe neurons at saccade onset among conditions during choice task in Figure S4.**

| ***Bad-pref GPe*** |  |  |  |  |  |  |
| --- | --- | --- | --- | --- | --- | --- |
| parametric bootstrap test (n = 10,000) | *p* |  |  |  |  |  |
| full model vs. null model | < .0001 |  |  |  |  |  |
| post-hoc  (pairwise t-test, Bonferroni correction) | Mean (SD) | Mean (SD) | *t* | *p* | 95% CI | effect size |
| **scene1** |  |  |  |  |  |  |
| (good, contra) vs (bad, contra) | –0.49 (1.11) | –1.69 (0.85) | 1.20 | < .0001 | [1.00 1.40] | 2.15 |
| (good, contra) vs (good, ipsi) | –0.49 (1.11) | –0.76 (1.07) | 0.28 | < .01 | [0.08 0.48] | 0.74 |
| (good, contra) vs (bad, ipsi) | –0.49 (1.11) | –1.66 (0.89) | 1.18 | < .0001 | [0.98 1.38] | 2.12 |
| (bad, contra) vs (good, ipsi) | –1.69 (0.85) | –0.76 (1.07) | –0.92 | < .0001 | [–1.12 –0.72] | –1.09 |
| (bad, contra) vs (bad, ipsi) | –1.69 (0.85) | –1.66 (0.89) | –0.02 | = .83 | [–0.22 0.18] | 0.28 |
| (good, ipsi) vs (bad, ipsi) | –0.76 (1.07) | –1.66 (0.89) | 0.90 | < .0001 | [0.70 1.10] | 1.68 |
| **scene2** |  |  |  |  |  |  |
| (good, contra) vs (bad, contra) | –0.70 (1.17) | –1.68 (0.86) | 0.98 | < .0001 | [0.78 1.18] | 1.84 |
| (good, contra) vs (good, ipsi) | –0.70 (1.17) | –0.90 (1.14) | 0.20 | < .05 | [0.00 0.41] | 0.62 |
| (good, contra) vs (bad, ipsi) | –0.70 (1.17) | –1.75 (0.87) | 1.06 | < .0001 | [0.86 1.26] | 1.93 |
| (bad, contra) vs (good, ipsi) | –1.68 (0.86) | –0.90 (1.14) | –0.78 | < .0001 | [–0.98 –0.58] | –0.88 |
| (bad, contra) vs (bad, ipsi) | –1.68 (0.86) | –1.75 (0.87) | 0.08 | = .45 | [–0.12 0.28] | 0.43 |
| (good, ipsi) vs (bad, ipsi) | –0.90 (1.14) | –1.75 (0.87) | 0.85 | < .0001 | [0.65 1.06] | 1.62 |
| **scene3** |  |  |  |  |  |  |
| (good, contra) vs (bad, contra) | –0.65 (1.12) | –1.66 (0.87) | 1.00 | < .0001 | [0.80 1.21] | 1.85 |
| (good, contra) vs (good, ipsi) | –0.65 (1.12) | –0.93 (1.13) | 0.28 | < .01 | [0.078 0.48] | 0.73 |
| (good, contra) vs (bad, ipsi) | –0.65 (1.12) | –1.64 (0.89) | 0.98 | < .0001 | [0.78 1.19] | 1.82 |
| (bad, contra) vs (good, ipsi) | –1.66 (0.87) | –0.93 (1.13) | –0.73 | < .0001 | [–0.93 –0.53] | –0.80 |
| (bad, contra) vs (bad, ipsi) | –1.66 (0.87) | –1.64 (0.89) | –0.02 | = .83 | [–0.22 0.18] | 0.27 |
| (good, ipsi) vs (bad, ipsi) | –0.93 (1.13) | –1.64 (0.89) | 0.71 | < .0001 | [0.50 0.91] | 1.39 |
| **scene4** |  |  |  |  |  |  |
| (good, contra) vs (bad, contra) | –0.58 (1.21) | –1.53 (0.89) | 0.95 | < .0001 | [0.75 1.15] | 1.77 |
| (good, contra) vs (good, ipsi) | –0.58 (1.21) | –0.83 (1.17) | 0.25 | < .05 | [0.05 0.45] | 0.69 |
| (good, contra) vs (bad, ipsi) | –0.58 (1.21) | –1.49 (0.91) | 0.91 | < .0001 | [0.71 1.11] | 1.71 |
| (bad, contra) vs (good, ipsi) | –1.53 (0.89) | –0.83 (1.17) | –0.70 | < .0001 | [–0.91 –0.50] | –0.77 |
| (bad, contra) vs (bad, ipsi) | –1.53 (0.89) | –1.49 (0.91) | –0.04 | = .68 | [–0.24 0.16] | 0.24 |
| (good, ipsi) vs (bad, ipsi) | –0.83 (1.17) | –1.49 (0.91) | 0.66 | < .0001 | [0.46 0.86] | 1.32 |

**Table S10. Summary of statistical test to compare the normalized neuronal activity of Bad-pref GPe neurons at saccade onset among conditions during choice task in Figure S4.**

| ***SNr*** |  |  |  |  |  |  |
| --- | --- | --- | --- | --- | --- | --- |
| parametric bootstrap test (n = 10,000) | *p* |  |  |  |  |  |
| full model vs. null model | < .0001 |  |  |  |  |  |
| post-hoc  (pairwise t-test, Bonferroni correction) | Mean (SD) | Mean (SD) | *t* | *p* | 95% CI | effect size |
| **scene1** |  |  |  |  |  |  |
| (good, contra) vs (bad, contra) | –1.06 (1.10) | 1.24 (1.20) | –2.30 | < .0001 | [–2.55 –2.06] | –2.63 |
| (good, contra) vs (good, ipsi) | –1.06 (1.10) | –0.79 (1.17) | –0.27 | < .05 | [–0.51 –0.02] | –0.31 |
| (good, contra) vs (bad, ipsi) | –1.06 (1.10) | 1.02 (1.33) | –0.28 | < .0001 | [–2.32 –1.83] | –2.37 |
| (bad, contra) vs (good, ipsi) | 1.24 (1.20) | –0.79 (1.17) | 2.04 | < .0001 | [1.79 2.28] | 2.32 |
| (bad, contra) vs (bad, ipsi) | 1.24 (1.20) | 1.02 (1.33) | 0.23 | = .071 | [–0.02 0.47] | 0.26 |
| (good, ipsi) vs (bad, ipsi) | –0.79 (1.17) | 1.02 (1.33) | –1.81 | < .0001 | [–2.05 –1.57] | –2.07 |
| **scene2** |  |  |  |  |  |  |
| (good, contra) vs (bad, contra) | –0.97 (1.09) | 1.45 (1.10) | –2.43 | < .0001 | [–2.67 –2.19] | –2.77 |
| (good, contra) vs (good, ipsi) | –0.97 (1.09) | –0.70 (1.13) | –0.27 | < .05 | [–0.52 –0.03] | –0.31 |
| (good, contra) vs (bad, ipsi) | –0.97 (1.09) | 1.32 (1.17) | –2.29 | < .0001 | [–2.53 –2.05] | –2.61 |
| (bad, contra) vs (good, ipsi) | 1.45 (1.10) | –0.70 (1.13) | 2.16 | < .0001 | [1.91 2.40] | 2.46 |
| (bad, contra) vs (bad, ipsi) | 1.45 (1.10) | 1.32 (1.17) | 0.14 | = .265 | [–0.11 0.38] | 0.16 |
| (good, ipsi) vs (bad, ipsi) | –0.70 (1.13) | 1.32 (1.17) | –2.02 | < .0001 | [–2.26 –1.77] | –2.30 |
| **scene3** |  |  |  |  |  |  |
| (good, contra) vs (bad, contra) | –1.28 (0.89) | 1.33 (1.18) | –2.61 | < .0001 | [–2.86 –2.37] | –2.98 |
| (good, contra) vs (good, ipsi) | –1.28 (0.89) | –0.91 (1.11) | –0.37 | < .01 | [–0.61 –0.12] | –0.42 |
| (good, contra) vs (bad, ipsi) | –1.28 (0.89) | 1.19 (1.08) | –2.47 | < .0001 | [–2.72 –2.23] | –2.82 |
| (bad, contra) vs (good, ipsi) | 1.33 (1.18) | –0.91 (1.11) | 2.25 | < .0001 | [2.00 2.49] | 2.56 |
| (bad, contra) vs (bad, ipsi) | 1.33 (1.18) | 1.19 (1.08) | 0.14 | = .259 | [–0.10 0.38] | 0.16 |
| (good, ipsi) vs (bad, ipsi) | –0.91 (1.11) | 1.19 (1.08) | –2.11 | < .0001 | [–2.35 –1.86] | –2.40 |
| **scene4** |  |  |  |  |  |  |
| (good, contra) vs (bad, contra) | –0.98 (1.18) | 1.39 (1.07) | –2.37 | < .0001 | [–2.61 –2.13] | –2.71 |
| (good, contra) vs (good, ipsi) | –0.98 (1.18) | –0.67 (1,18) | –0.31 | < .05 | [–0.55 –0.06] | –0.35 |
| (good, contra) vs (bad, ipsi) | –0.98 (1.18) | 1.22 (1.21) | –2.19 | < .0001 | [–2.44 –1.95] | –2.50 |
| (bad, contra) vs (good, ipsi) | 1.39 (1.07) | –0.67 (1,18) | 2.06 | < .0001 | [1.82 2.31] | 2.36 |
| (bad, contra) vs (bad, ipsi) | 1.39 (1.07) | 1.22 (1.21) | 0.18 | = .150 | [–0.06 0.42] | 0.20 |
| (good, ipsi) vs (bad, ipsi) | –0.67 (1,18) | 1.22 (1.21) | –1.89 | < .0001 | [–2.13 –1.64] | –2.15 |

**Table S11. Summary of statistical test to compare the normalized neuronal activity of SNr neurons at saccade onset among conditions during choice task in Figure S4.**

| ***Injection into GPe during choice task*** |  |  |  |  |  |  |
| --- | --- | --- | --- | --- | --- | --- |
| parametric bootstrap test (n = 10,000) | *p* |  |  |  |  |  |
| full model vs. null model | < .001 |  |  |  |  |  |
| post-hoc  (pairwise t-test, Bonferroni correction) | Mean (ms)  (SD) | Mean(ms)  (SD) | *t* | *p* | 95% CI | effect size |
| CPP+NBQX Contra Good pre vs. post | 195.89 (6.17) | 238.93  (15.99) | –0.19 | < .0001 | [–0.15 0.12] | –0.19 |
| Saline Contra Good pre vs. post | 192.81  (3.43) | 187.75  (8.93) | 0.02 | = .52 | [–0.04 0.08] | 0.02 |
| CPP+NBQX Contra Bad pre vs. post | 300.13  (23.16) | 317.00  (62.05) | –0.02 | = .41 | [–0.07 0.03] | –0.02 |
| Saline Contra Bad pre vs. post | 292.69  (16.09) | 285.19  (10.27) | 0.02 | = .48 | [–0.03 0.07] | 0.02 |
| CPP+NBQX Ipsi Good pre vs. post | 192.94  (10.29) | 189.63  (11.89) | 0.01 | = .76 | [–0.05 0.07] | 0.01 |
| Saline Ipsi Good pre vs. post | 193.69  (5.99) | 186.50  (7.22) | 0.04 | = .25 | [–0.03 0.10] | 0.04 |
| CPP+NBQX Ipsi Bad pre vs. post | 297.31  (32.50) | 295.13  (35.22) | 0.03 | = .32 | [–0.03 0.08] | 0.03 |
| Saline Ipsi Bad pre vs. post | 288.38  (28.57) | 280.69  (25.01) | 0.02 | = .37 | [–0.03 0.07] | 0.07 |
| ***Injection into SNr during choice task*** |  |  |  |  |  |  |
| parametric bootstrap test (n = 10,000) | *p* |  |  |  |  |  |
| full model vs. null model | < .001 |  |  |  |  |  |
| post hoc  (pairwise t-test, Bonferroni correction) | Mean (ms)  (SD) | Mean (ms)  (SD) | *t* | *p* | 95% CI | effect size |
| CPP+NBQX Contra Good pre vs. post | 197.06 (5.47) | 172.81  (13.36) | 0.13 | < .001 | [0.06 0.21] | 0.13 |
| Saline Contra Good pre vs. post | 193.19  (5.25) | 187.94  (3.49) | 0.03 | = .45 | [–0.04 0.10] | 0.03 |
| CPP+NBQX Contra Bad pre vs. post | 280.06  (14.10) | 181.31  (21.82) | 0.43 | < .0001 | [0.35 0.52] | 0.43 |
| Saline Contra Bad pre vs. post | 274.19  (17.74) | 279.13  (15.53) | –0.02 | = .55 | [–0.08 0.04] | –0.02 |
| CPP+NBQX Ipsi Good pre vs. post | 198.75  (5.66) | 213.06  (15.50) | –0.07 | = .05 | [–0.14 –0.01] | –0.07 |
| Saline Ipsi Good pre vs. post | 196.63  (3.92) | 195.13  (5.59) | 0.01 | = .82 | [–0.06 0.08] | 0.01 |
| CPP+NBQX Ipsi Bad pre vs. post | 296.69  (28.17) | 287.31  (66.49) | 0.03 | = .27 | [–0.03 0.09] | 0.03 |
| Saline Ipsi Bad pre vs. post | 303.19  (23.87) | 287.13  (33.66) | 0.05 | = .06 | [–0.01 0.11] | 0.05 |

**Table S12. Summary of statistical test to compare the effects of CPP + NBQX injection into GPe and SNr during choice task in Figure 3.**

| ***Injection into GPe during choice task*** |  |  |  |  |  |  |
| --- | --- | --- | --- | --- | --- | --- |
| Accept Bad object |  |  |  |  |  |  |
| parametric bootstrap test (n = 10,000) | *p* |  |  |  |  |  |
| full model vs. null model | = 1.00 |  |  |  |  |  |
| Return for Bad object |  |  |  |  |  |  |
| parametric bootstrap test (n = 10,000) | *p* |  |  |  |  |  |
| full model vs. null model | < .001 |  |  |  |  |  |
| post-hoc  (pairwise t-test, Bonferroni correction) | Mean (%)  (SD) | Mean (%) (SD) | *t* | *p* | 95% CI | effect size |
| CPP+NBQX  Return Contra Bad pre vs. post | 68.63  (10.94) | 47.40  (42.99) | 1.05 | < .0001 | [0.76 1.34] | 1.05 |
| Saline  Return Contra Good pre vs. post | 70.11  (12.67) | 57.16  (22.66) | –0.20 | = .22 | [–0.52 0.12] | –0.02 |
| CPP+NBQX  Return Ipsi Bad pre vs. post | 77.30  (14.61) | 80.52  (6.44) | 0.61 | < .0001 | [0.32 0.89] | 0.61 |
| Saline  Return Ipsi Bad pre vs. post | 74.65  (15.14) | 78.42  (11.26) | –0.25 | = .14 | [–0.56 0.06] | –0.25 |
| Stay for Bad object |  |  |  |  |  |  |
| parametric bootstrap test (n = 10,000) | *p* |  |  |  |  |  |
| full model vs. null model | < .001 |  |  |  |  |  |
| post-hoc  (pairwise t-test, Bonferroni correction) | Mean (%)  (SD) | Mean (%) (SD) | *t* | *p* | 95% C.I. | effect size |
| CPP+NBQX  Return Contra Bad pre vs. post | 29.89  (13.79) | 46.54  (37.18) | –0.75 | < .0001 | [–1.03 –0.47] | –0.75 |
| Saline  Return Contra Good pre vs. post | 29.35  (13.17) | 39.33  (24.33) | 0.20 | = .22 | [–0.78 –0.19] | 0.20 |
| CPP+NBQX  Return Ipsi Bad pre vs. post | 22.70  (14.61) | 19.00  (6.77) | –0.49 | < .0001 | [–0.12 0.53] | –0.49 |
| Saline  Return Ipsi Bad pre vs. post | 25.35  (15.14) | 21.58  (11.26) | 0.24 | = .14 | [–0.08 0.55] | 0.24 |

**Table S13. Summary of statistical test to compare the effects of CPP + NBQX injection into GPe while monkeys chose actions for Bad object during choice task in Figure S10.**

| ***Injection into SNr during Choice task*** |  |  |  |  |  |  |
| --- | --- | --- | --- | --- | --- | --- |
| Accept Bad object |  |  |  |  |  |  |
| parametric bootstrap test (n = 10,000) | *p* |  |  |  |  |  |
| full model vs. null model | = 1.00 |  |  |  |  |  |
| Return for Bad object |  |  |  |  |  |  |
| parametric bootstrap test (n = 10,000) | *p* |  |  |  |  |  |
| full model vs. null model | < .001 |  |  |  |  |  |
| post-hoc  (pairwise t-test, Bonferroni correction) | Mean (%)  (SD) | Mean (%) (SD) | *t* | *p* | 95% CI | effect size |
| CPP+NBQX  Return Contra Bad pre vs. post | 77.88  (15.25) | 86.32  (12.39) | –0.57 | < .0001 | [–1.00 –0.14] | –0.57 |
| Saline  Return Contra Good pre vs. post | 74.57  (12.37) | 71.45  (12.26) | 0.22 | = .22 | [–0.10 0.55] | 0.22 |
| CPP+NBQX  Return Ipsi Bad pre vs. post | 64.19  (20.23) | 41.23  (23.41) | 1.06 | < .0001 | [0.72 1.41] | 1.06 |
| Saline  Return Ipsi Bad pre vs. post | 60.42  (16.09) | 62.20  (18.07) | 0.07 | = .14 | [–0.23 0.36] | 0.07 |
| Stay for Bad object |  |  |  |  |  |  |
| parametric bootstrap test (n = 10,000) | *p* |  |  |  |  |  |
| full model vs. null model | < .001 |  |  |  |  |  |
| post-hoc  (pairwise t-test, Bonferroni correction) | Mean (%)  (SD) | Mean (%) (S.D.) | *t* | *p* | 95% CI | effect size |
| CPP+NBQX  Return Contra Bad pre vs. post | 22.12  (15.25) | 5.55  (10.23) | 1.30 | < .0001 | [0.79 1.82] | 1.30 |
| Saline  Return Contra Good pre vs. post | 25.43  (12.37) | 28.05  (11.96) | –0.18 | = .22 | [–1.38 –0.69] | –0.18 |
| CPP+NBQX  Return Ipsi Bad pre vs. post | 35.53  (20.35) | 57.50  (21.99) | –1.03 | < .0001 | [–0.50 0.15] | –1.03 |
| Saline  Return Ipsi Bad pre vs. post | 39.46  (16.19) | 37.56  (17.80) | –0.04 | = .14 | [–0.34 0.26] | –0.04 |

**Table S14. Summary of statistical test to compare the effects of CPP + NBQX injection into SNr while monkeys chose actions for Bad object during Choice task in Figure S10.**

**Table S15. Summary of statistical test to compare the effects of CPP + NBQX injection into GPe and *SNr during fixation task in Figure 3.***

| ***Injection into GPe during fixation task*** |  |  |  |  |  |  |
| --- | --- | --- | --- | --- | --- | --- |
| parametric bootstrap test (n = 10,000) | *p* |  |  |  |  |  |
| full model vs. null model | = .05 |  |  |  |  |  |
| ***Injection into SNr during fixation task*** |  |  |  |  |  |  |
| parametric bootstrap test (n = 10,000) | *p* |  |  |  |  |  |
| full model vs. null model | < .01 |  |  |  |  |  |
| post-hoc  (pairwise t-test, Bonferroni correction) | Mean (%)  (SD) | Mean (%) (SD) | *t* | *p* | 95% CI | effect size |
| CPP+NBQX Contra Good pre vs. post | 1.33  (2.30) | 63.91  (26.74) | –5.29 | < .0001 | [–6.46 –4.12] | –5.29 |
| Saline Contra Good pre vs. post | 1.47  (1.49) | 3.05  (3.36) | –0.59 | = .26 | [–1.61 0.44] | –0.59 |
| CPP+NBQX Contra Bad pre vs. post | 2.40  (2.59) | 55.93  (24.27) | –4.11 | < .0001 | [–5.02 –3.20] | –4.11 |
| Saline Contra Bad pre vs. post | 1.40  (2.07) | 1.13  (1.69) | 0.03 | = .96 | [–1.29 1.35] | 0.03 |
| CPP+NBQX Ipsi Good pre vs. post | 2.09  (3.50) | 0.25  (6.61) | 1.74 | = .10 | [–0.35 3.84] | 1.74 |
| Saline Ipsi Good pre vs. post | 0.40  (0.71) | 0.63  (1.10) | –0.20 | = .84 | [–2.16 1.76] | –0.2 |
| CPP+NBQX Ipsi Bad pre vs. post | 1.51  (2.78) | 1.20  (2.23) | 0.2 | = .75 | [–1.04 1.45] | 0.2 |
| Saline Ipsi Bad pre vs. post | 0.36  (0.94) | 0.33  (0.87) | –0.05 | = .97 | [–2.82 2.72] | –0.05 |
